# Controllable Gap Junctions by Vitamin B_12_ and Light

**DOI:** 10.1101/2025.05.28.656405

**Authors:** Duo Cui, Shuzhang Liu, Xinyu Huang, Xiaohan Alex Tang, Min Zheng, Zonglin He, Rui Xu, Chenbo Sun, Yingjie Xu, Renjun Tu, Peng Zou, Ting Xie, Fei Sun

**Affiliations:** Department of Chemical and Biological Engineering, The Hong Kong University of Science and Technology, Clear Water Bay, Kowloon, Hong Kong SAR, China; Division of Life Science, The Hong Kong University of Science and Technology, Clear Water Bay, Kowloon, Hong Kong SAR, China; College of Chemistry and Molecular Engineering, Synthetic and Functional Biomolecules Center, Beijing National Laboratory for Molecular Sciences, Key Laboratory of Bioorganic Chemistry and Molecular Engineering of the Ministry of Education, Peking University, Beijing, China; School of Life Science and Technology, Southeast University, Nanjing, China; Biomedical Research Institute, Shenzhen Peking University–The Hong Kong University of Science and Technology Medical Center, Shenzhen 518036, China; Shenzhen Bay Laboratory, Shenzhen 518036, China

**Author notes:** These authors contributed equally to this work (Duo Cui, Shuzhang Liu, Xinyu Huang).

## Abstract

Gap junctions, mediate rapid signal transduction between contiguous cells, which are indispensable for multicellular organisms to coordinate cellular activities across numerous physiological processes. However, precise control of gap junctions remains elusive. Herein, we present CarGAP, a single-component chemo-optogenetic tool that utilizes the C-terminal adenosylcobalamin (AdoB_12_) binding domain of a photoreceptor protein (i.e., CarH_C_) to achieve reversible control over both vertebrate and invertebrate gap junctions with spatiotemporal precision. The vertebrate CarGAP (i.e., Cx-CarGAP), created by genetically fusing connexins with CarH_C_ in mammalian cells, can efficiently block the gap junction channels through AdoB_12_-induced protein oligomerization, and subsequently reinstate them via green light-induced protein disassembly. We further introduced the CarGAP system (i.e., Inx-CarGAP) to the *Drosophila* ovary, enabling reversible control over the heterotypic gap junctions formed by innexin2 (Inx2) and innexin4 (Inx4, also known as zero population growth, Zpg), thereby uncovering the roles of gap junctions in stem cell-niche interactions. This study illustrates CarGAP as a generalizable chemo-optogenetic tool for interrogating the functions of gap junctions in various biological contexts.

## Introduction

Gap junctions are ubiquitous intercellular channels found in virtually all tissues, which intricately connect an array of cell types and facilitate speedy intercellular communications^1^. In vertebrates, gap junctions connect adjacent cells via hexameric transmembrane proteins known as connexins, while their counterparts in invertebrate cells use octameric protein complexes known as innexins^2^. The evolutionary relationship between connexins and innexins remains enigmatic, given their low sequence homology but high functional similarity^3^. The nanometer-sized channels formed by, gap junction channels in adjacent cells allow for their direct exchange of ions^4^, second messengers^5–7^, microRNAs^8^, amino acids^9^, and oligopeptides^10^, thus impacting a wide range of physiological processes, such as embryonic development^5^, immune function^6^, cardiac contraction^11^, and neuronal synchronization^12, 13^. Malfunction of gap junctions has been implicated in various diseases, ranging from cardiovascular disorders^11^ to developmental abnormalities^14^ and cancer progression^15^.

Gap junctions are dynamically regulated under various physiological conditions, ensuring continuous intercellular communication and synchronization between adjacent cells. They may close in response to certain stimuli such as elevated calcium levels^16^, abnormal pH^17^, or posttranslational modifications^18^. In developmental biology, gap junctions are highly expressed in early developmental stages, mediating secondary messenger signaling^19^. Traditional genetic methods such as gene knockout/knockdown and loss-of-function mutations, though commonly used, permanently and indiscriminately abolish all functions of gap junction proteins. This fails to differentiate those channel-dependent roles from channel-independent ones and often results in severe consequences such as embryonic lethality^20, 21^. In neurobiology, there are two main modalities of synaptic transmission: chemical synapses and electrical synapses, the latter being mediated by gap junctional channels (GJCs)^22^. Molecular tools have been disproportionately developed for controlling chemical synapses, but not for GJC-mediated electrical synpases^23–25^.

Commonly used small-molecule inhibitors, such as weak organic acids and strong reducing reagents, only partially impair GJC conductance through intracellular acidification and disulfide bond cleavage, respectively. However, these inhibitors often exert adverse nonspecific effects on organisms, limiting their broader applications^26, 27^. Peptide-based blockers have been designed to target the loop region of gap junction proteins, offering enhanced specificity and biocompatibility, but they suffer from poor pharmacokinetics due to rapid degradation in vivo^28^. Chromophore-assisted light inactivation (CALI) technology has been used to inactivate gap junctions via reactive oxygen species photochemically generated by either (1) genetically encoded EGFP (light intensity: 1.3 MW/cm²)^29^ or (2) a synthetic membrane-permeable, red biarsenical dye bound to tetra-cysteine motifs (light intensity: 17 W/cm²)^30^. Both CALI methods require relatively high light intensities^30, 31^. While photocleavable PhoCl has been successfully employed for selective activation of pannexin hemichannels in a pioneering study^32^, the development of precise and more gentle methods to control canonical connexin- or innexin-based GJCs - the critical mediators of direct cell-to-cell communication in multicellular organisms - remains an outstanding challenge.

Recently, the vitamin B_12_-dependent photoreceptor protein CarH has emerged as a robust chemo-optogenetic tool for controlling protein assembly and disassembly in vitro^33, 34^, as well as gene expression in vivo^35, 36^. Originally identified as a transcriptional regulator of carotenoid biosynthesis in *Thermus thermophilus*^37, 38^, CarH consists of an N-terminal DNA-binding domain and a C-terminal adenosylcobalamin (AdoB_12_)-binding domain (CarH_C_). In its apo state, CarH_C_ is monomeric but undergoes tetramerization upon AdoB_12_ binding in the dark. Subsequent green-light illumination triggers the disassembly of tetramers into monomers, accompanied by the cleavage of C-Co, adenosyl release, and formation of bis-His-ligated B ^39^. Due to the high photosensitivity of AdoB_12_, even low-intensity green light (μW/cm² range) efficiently drives the protein disassembly, pointing to CarH_C_’s potential as a gentle and precise tool for optogenetics^35, 40^. In this study, by leveraging CarH_C_, we developed CarGAP, a single-component chemo-optogenetic tool for precise and reversible control of both vertebrate (connexin-based) and invertebrate (innexin-based) GJCs. Using a connexin-based CarGAP, we demonstrated AdoB_12_- and light-dependent regulation of intercellular mass transfer (e.g., fluorescent dyes and 2’,3’-cGAMP) and electrical coupling in mammalian cells. Furthermore, innexin-based CarGAPs enabled in vivo investigation of cAMP transport and signaling through heterotypic GJCs in the *Drosophila* germarium, revealing molecular-level insights into stem cell–niche interactions. Given the prevalence and diversity of GJCs, CarGAP offers a powerful approach to dissect their functions in various biological contexts.

## Results

### Design and functional validation of CarGAP in mammalian cells

To demonstrate the feasibility of creating chemo-optogenetically controllable gap junctions in living cells, we selected well-characterized connexin 43 (Cx43)^41, 42^, the most abundant connexin isoform in vertebrates. As previous studies have shown that placing fluorescent proteins at the C-terminus of Cx43 does not compromise its channel-dependent functionality^43, 44^, we created CarGAP by genetically fusing CarH_C_ at the C-terminus of human Cx43 (GJA1) as well. We conjectured that the possible formation of a dimer-dimers type of tetramer and a dimer upon six CarH_C_ monomers binding to AdoB_12_ could block the gap junction channel from the intracellular side, thus halting cell-to-cell material exchanges and communications, which could be reversed by CarH_C_ disassembly induced by green light irradiation, a process that is accompanied with the C-Co cleavage of AdoB_12_ and the bis-His-Co ligation (Fig. 1a,b and Supplementary Fig. 1). Neuro-2a (N2A) cells, a fast-growing mouse neuroblastoma cell line, reportedly show no detectable connexin-based GJCs and therefore can serve as an ideal cellular system for studying recombinant GJCs^45, 46^. Most human connexins, including Cx43, have been functionally expressed in N2A cells^47, 48^. The N2A cells were genetically engineered to stably express CarGAP via the pLVX-Puro lentiviral expression vector (Fig. 1c). To assess the channel function of CarGAP, we performed a dye transfer assay between co-cultured CarGAP N2A cells. Donor cells were labeled with two fluorescent dyes, a red- (DiD) or orange-fluorescent (Dil) lipophilic tracer for plasma membranes and green-fluorescent calcein AM for cytoplasmic staining, and then mixed and co-cultured with unlabeled CarGAP N2A recipient cells at a ∼1:40 donor-to-recipient ratio, following an established protocol^49, 50^. The membrane-bound nontransferable DiD served to differentiate the donor cells from the recipient cells, while the hydrophobic calcein AM could be hydrolyzed into hydrophilic calcein molecule upon entering the cytoplasm, of which the transferability from the donor cells to the co-cultured recipient cells is highly indicative of the function of GJCs (Fig. 1d).

**Fig. 1.**
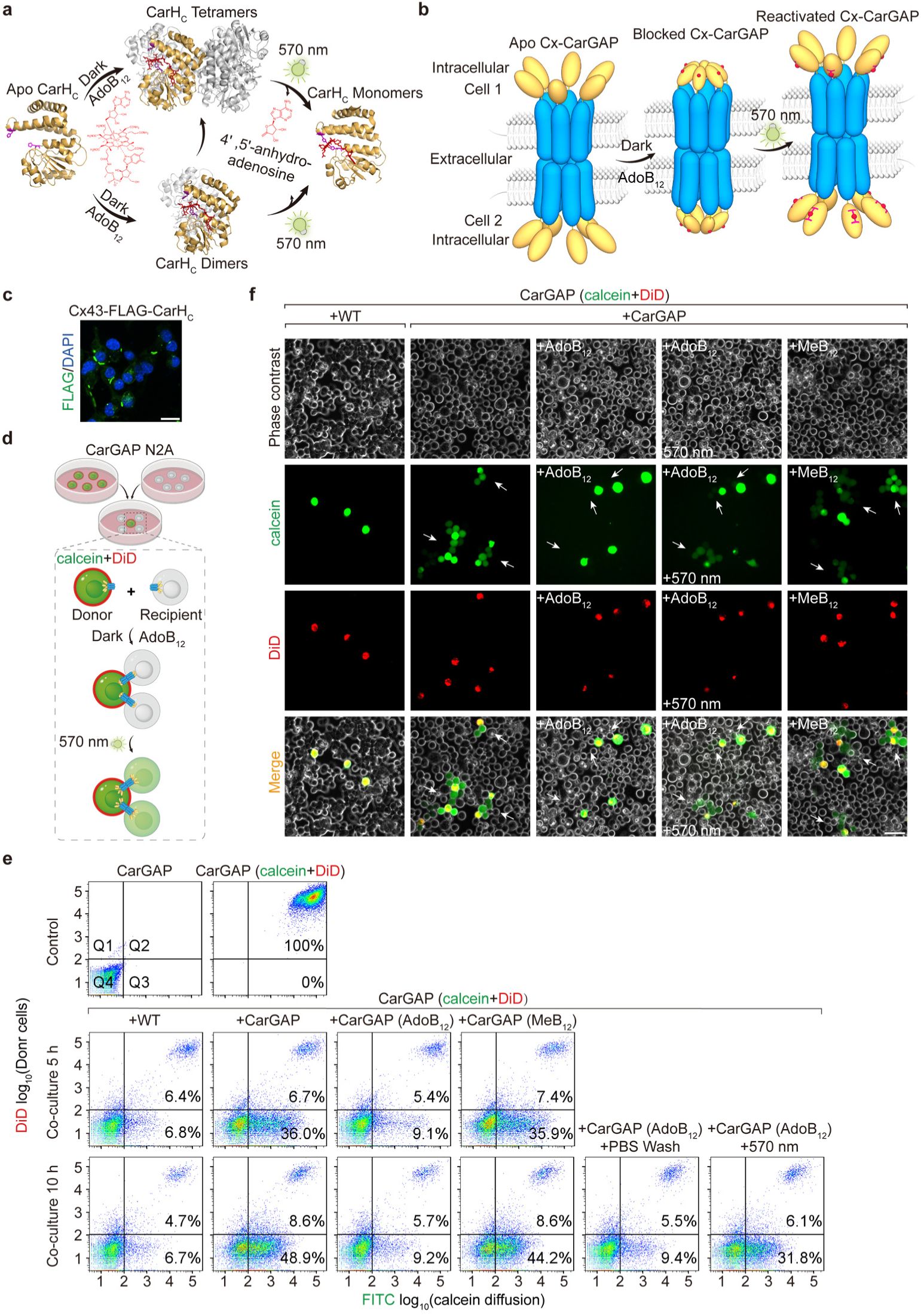
Chemo-optogenetically controlled intercellular transfer of fluorescent dyes via connexin-CarGAP. **a**, Photochemistry of CarH_C_. CarH_C_ monomers bind to AdoB_12_ in the dark, forming dimers or tetramers (PDB: 5C8A), which subsequently disassemble into monomers (PDB: 5C8F) upon exposure to 570-nm green light, accompanied by the photolysis of AdoB_12_ and the release of 4′,5′-anhydroadenosine. **b**, Schematic representation of Connexin-CarGAP (Cx-CarGAP). CarGAP functions as a molecular valve between adjacent cells. In the dark, AdoB_12_ induces the oligomerization of CarH_C_, blocking the connexin gap junction channel. Upon illumination with 570-nm green light, CarGAP restores its channel-dependent function. **c**, Immunofluorescence image of Cx-CarGAP. N2A cells stably transfected with *Cx43-FLAG-carH_C_* are shown. Scale bar: 20 µm. Detailed descriptions of the genetic constructs are provided in Supplementary Table 1. **d**, Schematic representation of the co-culturing experimental procedure for assessing Cx-CarGAP. Donor cells are first labeled with the membrane-bound dye DiD (red) and the permeable dye calcein AM (green), then co-cultured with unlabeled recipient cells at a ratio of ∼1:40 (∼1:20) and a total density of ∼50,000 cells/cm². The impermeable DiD differentiates donor from recipient cells, while the transfer of calcein from donor to recipient cells indicates the functionality of the gap junction channels. AdoB_12_ is added under dark conditions to examine the closure of Cx-CarGAP, as indicated by the absence of calcein (green fluorescence) in recipient cells surrounding the donor cell. After co-culturing in the presence of AdoB_12_ for 5 hours, the cells are exposed to green light (570 nm; 10 mW/cm^2^) for 5 minutes to assess the reopening of the CarGAP channel *in situ*, as evidenced by the transfer of calcein from donor cells to adjacent cells. **e**, Flow cytometry analysis of dye transfer between N2A cells to assess the function of GJCs. FACS plots are gated into 4 quadrants (Q1, Q2, Q3, Q4). Donor and recipient cells were mixed at a ratio of ∼1:20 and a density of ∼50,000 cells/cm², then co-cultured for 5 and 10 hours. Donor cells, characterized by both red (DiD) and green fluorescence (calcein), are located in quadrant Q2, while recipient cells, characterized by green fluorescence (calcein) only, are located in quadrant Q3 of the FACS plots. The corresponding histogram plots of FITC are shown in Supplementary Fig. 4. All experiments were performed at least three times to ensure consistency. **f**, Fluorescence images showing the transfer of calcein AM from donor cells to recipient cells under various conditions: wild-type N2A to wild-type N2A, CarGAP N2A to CarGAP N2A, CarGAP N2A to CarGAP N2A with AdoB_12_, CarGAP N2A to CarGAP N2A with AdoB_12_ followed by light illumination (570 nm; 5 min), and CarGAP N2A to CarGAP N2A with MeB_12_. Red-channel images (633 nm) show donor cells pre-labeled with DiD, while green-channel images (488 nm) display donor cells pre-labeled with calcein AM, as well as recipient cells that acquire calcein through functioning gap junction channels. Transfer of calcein occurred between CarGAP N2A cells, including those in the presence of MeB_12_ (a vitamin B_12_ variant that does not induce protein oligomerization), but were blocked in the presence of AdoB_12_. This blockage was abolished by light irradiation. Wild-type N2A cells, which lack intrinsic connexin gap junctions and disallow intercellular transfer of calcein, served as controls. Scale bar: 50 µm. All experiments were performed at least three times to ensure consistency.

To determine the speed of GJC formation in co-cultured cells, we initially conducted time-course measurements of intercellular dye transfer. Flow cytometry analysis of co-cultured CarGAP N2A cells revealed little calcein transfer at 3 hours post-mixing, suggesting insufficient time for reforming GJCs (Supplementary Fig. 2). By contrast, intercellular calcein diffusion became apparent after 5 hours of co-culturing, with CarGAP N2A cells adjacent to prelabeled cells displaying clear green fluorescence (Supplementary Fig. 2 and Fig. 1e), which thus established 5 hours as the optimal timepoint for subsequent functional validation of CarGAP. Wild-type N2A cells showed no dye transfer even after 5 hours, consistent with their inherent gap junction deficiency.

To quantitatively evaluate CarGAP-mediated calcein transfer under various conditions, we performed flow cytometry analyses using cells co-cultured at a donor-to-recipient ratio of ∼1:20. In the absence of AdoB_12_, calcein transfer was observed in 36.0% of CarGAP N2A cells after 5 hours of co-culturing, increasing to 48.9% by 10 hours (Q3 population, green fluorescence only; Fig. 1e). AdoB_12_ – actively uptaken by cells through receptor-mediated endocytosis^51, 52^ – inhibited calcein transfer in CarGAP N2A cells in a dose-dependent manner, with 50 μM AdoB_12_ causing significant suppression and 500 μM achieving almost complete blockage within 5 hours (Supplementary Fig. 3). AdoB_12_ (500 μM) further reduced calcein-positive CarGAP N2A cells to just 9.2% (Q3 population) after 10 hours of co-culturing. This inhibition reached near-baseline levels observed in gap junction-deficient wild-type N2A cells (6.7%) (Fig. 1e). The AdoB_12_-induced inhibition persisted under dark conditions, remaining effective even after phosphate-buffered saline (PBS) washing and an additional 5 hours of co-culturing. Nevertheless, brief green light exposure (5 min) completely reversed this blockage, restoring transfer efficiency to 31.8% of recipient cells in Q3. In contrast, MeB_12_, a vitamin B_12_ variant that binds to CarH_C_ without inducing oligomerization^53^, showed minimal effect on calcein transfer, with 35.9% (5 h) and 44.2% (10 h) of recipient cells remaining in Q3, suggesting that CarH_C_ oligomerization is required for channel inhibition.

Complementing our flow cytometry data, fluorescence microscopy confirmed complete suppression of calcein transfer in AdoB_12_-treated CarGAP N2A cells under dark conditions, demonstrating that AdoB_12_-induced CarH_C_ oligomerization effectively blocks channel function. This inhibition can be readily reversed, with 5-minute green light exposure (10 mW/cm²) fully restoring intercellular calcein diffusion (Fig. 1f). Once again MeB_12_ - which cannot induce CarH_C_ oligomerization - failed to inhibit dye transfer (Fig. 1f), further establishing oligomerization as the essential mechanism for channel blockage. Notably, AdoB_12_ concentrations up to ∼500 μM maintained excellent cytocompatibility, consistent with known vitamin B_12_ safety profiles^54^.

Collectively, these results establish connexin-based CarGAP as a robust chemo-optogenetic platform for reversible regulation of GJC-mediated molecular transfer in mammalian cells.

### Chemo-optogenetically controlled transport of 2’3’-cGAMP across mammalian cells

The second messenger, 2’3’-cyclic GMP-AMP (2’3’-cGAMP), plays a pivotal role in innate immunity and viral defense^55^. In a canonical process, 2’3’-cGAMP, synthesized by cyclic GMP-AMP synthase (cGAS) in response to cytosolic double-stranded DNA, activates the stimulator of interferon genes (STING) signaling pathway within the producing cells, which further triggers a cascade of antiviral responses in a type I interferon-dependent manner^55^. Alternatively, in a process known as bystander immunity^6^, 2’3’-cGAMP transfers from producing cells to adjacent cells through connexin gap junctions, thus promoting STING signaling and antiviral immunity in the abutted cells independently of type I IFN signaling. The activation of STING by cGAMP induces its oligomerization into a supramolecular complex and prompts its translocation from the endoplasmic reticulum (ER) to a perinuclear compartment in Gogi, which can be readily observed using fluorescence microscopy^56^ (Fig. 2a). Native N2A cells, while lacking intrinsic Cx43 and STING, possess endogenous cGAS capable of sensing and responding to cytosolic double-stranded DNA (dsDNA), such as plasmids. This characteristic hampers their utility as a model cell line for the transient transfection of *STING*, as it can lead to autoactivation following transfection (Supplementary Fig. 5). In contrast, native HEK293T cells do not express either cGAS or STING^57^ but possess abundant endogenous gap junctions, including Cx43. Moreover, co-culturing the two cell lines indeed formed functional gap junctions, as evidenced by the pronounced calcein transfer from CarGAP N2A cells to neighboring HEK293T cells (Fig. 2b). The complementarity between N2A cells and HEK293T cells renders them ideal donors and recipients, respectively, to examine the possibility of controlling intercellular transport of 2’3’-cGAMP. To accomplish this, we boosted the synthesis of cGAMP in both WT and CarGAP N2A cells by overexpressing cGAS, along with HT-DNA activation, followed by mixing and co-culturing with the recipient HEK293T cells transfected with *STING-egfp* at a ratio of ∼1:40 (Fig. 2c).

**Fig. 2.**
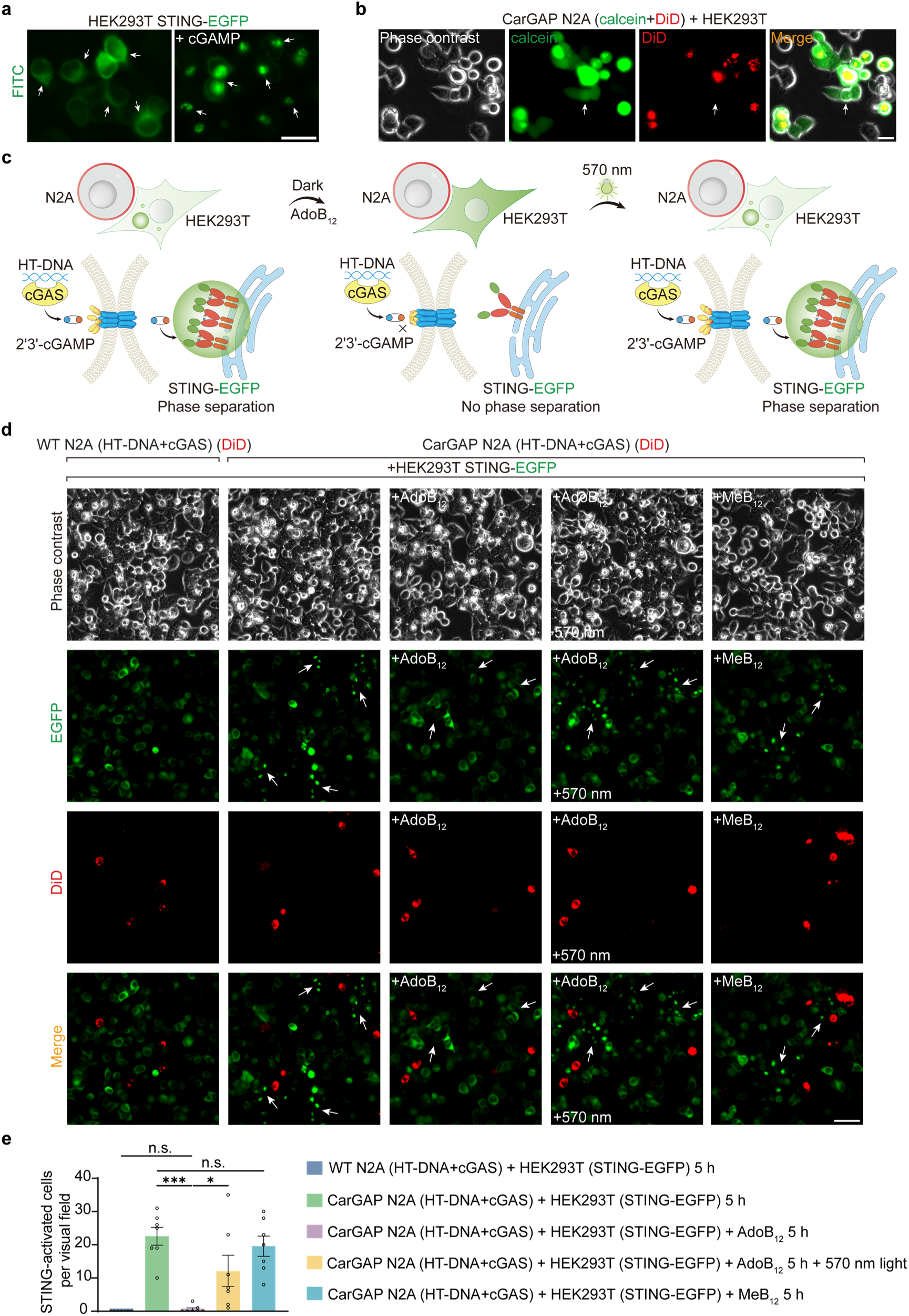
Chemo-optogenetically controlled intercellular transport of 2′3′-cGAMP via connexin-CarGAP. **a**, Exogenous cGAMP induces the oligomerization and translocation of STING-EGFP in HEK293T cells, which lack endogenous cGAS to produce 2′3′-cGAMP. Scale bar: 20 µm. **b**, CarGAP-mediated transport of calcein from CarGAP N2A cells to adjacent HEK293T cells. Scale bar: 20 µm. **c**, Schematic representation of chemo-optogenetically controlled intercellular cGAMP transport enabled by CarGAP. The closure of CarGAP induced by AdoB_12_ blocks the transfer of cGAMP from N2A to HEK293T cells, which is subsequently restored upon exposure to green light. **d**, Co-culturing with N2A cells pre-treated with HT-DNA (5 μg/ml) induces LLPS of STING-EGFP in HEK293T cells via CarGAP. Both wild-type and CarGAP N2A cells express endogenous cGAS but lack native connexin gap junctions. HEK293T cells are deficient in native cGAS production. N2A cells overexpressing cGAS and pre-labeled with DiD were co-cultured with STING-EGFP HEK293T cells for 5 hours under various conditions: without B_12_ (none), with 500 μM AdoB_12_, with 500 μM AdoB_12_ followed by 5 minutes of green light exposure (10 mW/cm²) and an additional 5-hour culture, and with 500 μM MeB_12_. Representative images are shown. Scale bar: 50 µm. **e**, Quantification of STING-activated HEK293T cells surrounding N2A donors. Data are presented as mean ± SEM (n = 7 different visual fields). A two-sided t-test was used. * p < 0.05; *** p < 0.001; n.s., not significant.

Contrary to WT N2A cells, which failed to elicit any bystander STING-EGFP translocation from ER to Gogi in adjacent HEK293T cells owing to the absence of functional gap junctions and intercellular cGAMP transport, CarGAP N2A cells were surrounded by HEK293T cells that exhibited clear STING-EGFP translocation, indicative of functional gap junction channels that mediated the transport of cGAMP from the producing CarGAP N2A cells to the nearby HEK293T cells. The intercellular transport of cGAMP was inhibited by AdoB_12_, as evidenced by few HEK293T cells with STING-EGFP translocation in the co-cultures in the presence of AdoB_12_ in the dark, strongly suggesting the blockage of CarGAP channels. The blockage of cGAMP transport from N2A cells to HEK293T cells was eliminated following a 5-minute green light irradiation, leading to considerable STING-EGFP translocation in the HEK293T cells (Fig. 2d,e). Once again, CarH_C_ oligomerization proved to be essential for the blockage of CarGAP channels and thus the transport of cGAMP, as MeB_12_, which binds to CarH_C_ but induces no oligomerization, exerted almost no effect on STING-EGFP in HEK293T, of which translocation remained. These findings not only corroborated the critical roles of gap junctions in cGAS-STING signaling and bystander immunity, but also highlighted the applicability of using CarGAP to manipulate the intercellular transport of crucial second messengers.

### Chemo-optogenetic control and far-red fluorescent recording of electrical coupling among adjacent cells

Gap junctions enable the instantaneous and reciprocal spread of current among adjacent cells^22^. The traditional patch-clamp technique has often been used to study electrophysiology in individual cells. However, applying this method to multiple cells connected through gap junctions remains challenging. It is therefore highly desirable to have a technique that not only enables precise control over the electrical coupling and dynamics within cell clusters but also facilitates straightforward recording of these events. Regarding the control of electrical coupling, CarGAP has provided a chemo-optogenetic way to modulate gap junction channels and therefore holds great promise for controlling electrical coupling across multiple cells. In terms of recording electrical dynamics, a series of hybrid fluorescent membrane voltage indicators have recently been created via the site-specific modification of rhodopsin proteins with synthetic small-molecule fluorophores. These indicators have enabled the optical mapping of gap junction-mediated electrical couplings across a monolayer of HEK293T cells^58^. We envisioned that combining the chemo-optogenetic tool CarGAP with the hybrid voltage indicators could lead to a methodology capable of chemo-optical control while also optically recording the electrical dynamics in an interconnected multicellular system. To avoid interfering with CarGAP, which is sensitive to green light, we selected a far-red hybrid voltage indicator, NAVI-Cy5 (unpublished), that responds to membrane voltage changes via the electrochromic Förster resonance energy transfer (eFRET) mechanism^59^ (Fig. 3a).

**Fig. 3.**
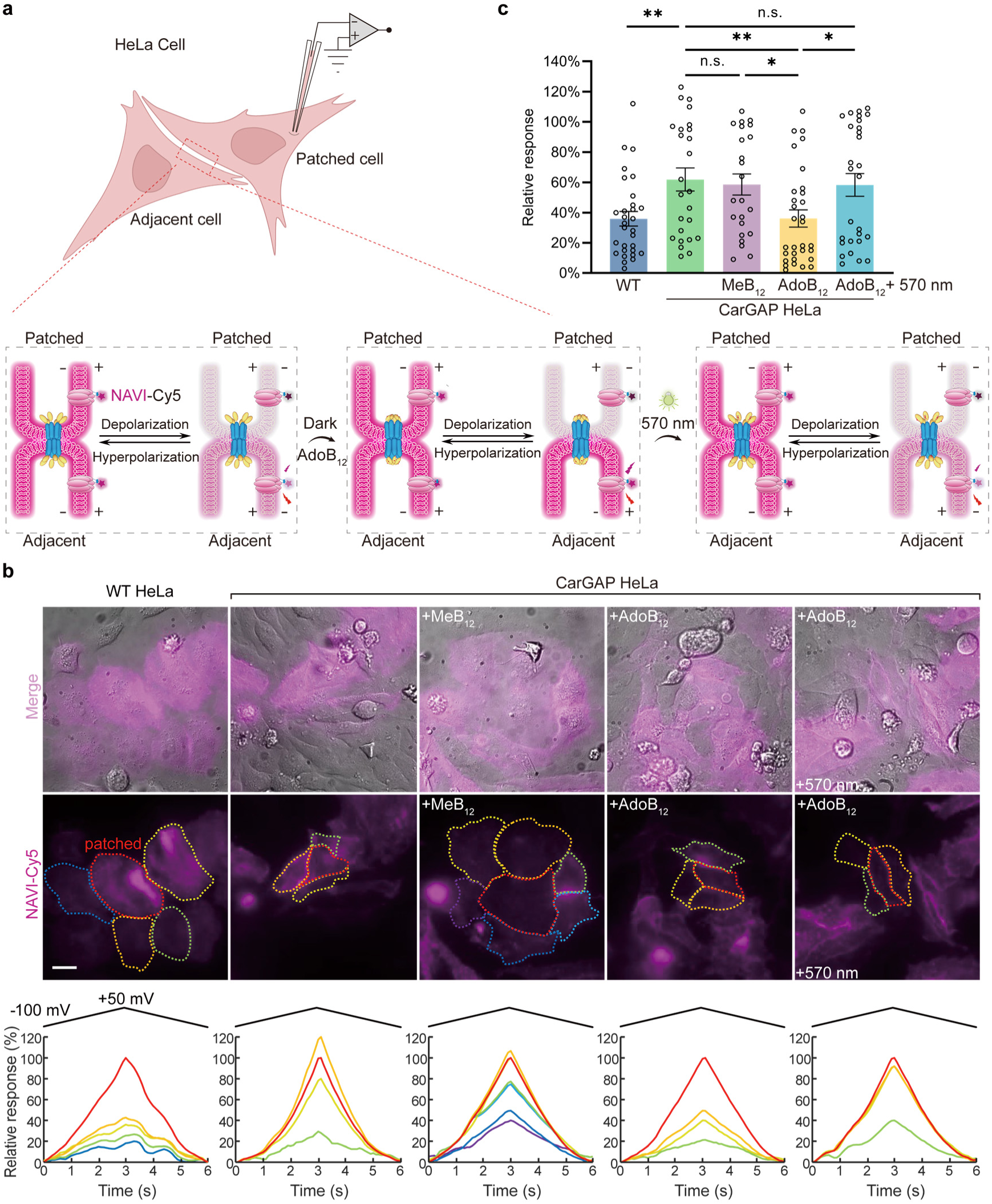
Chemo-optogenetically controlled electrical coupling between cells via CarGAP. **a**, Schematic illustration of controlled electrical coupling between HeLa cells enabled by CarGAP, visualized using the fluorescent voltage indicator NAVI-Cy5. NAVI-Cy5 detects changes in membrane potential through an electrochromic FRET mechanism; membrane depolarization results in decreased Cy5 fluorescence emission, while hyperpolarization leads to increased emission. AdoB_12_-induced protein oligomerization blocks CarGAP, electrically uncoupling adjacent cells, whereas subsequent light-induced disassembly restores electrical coupling. **b**, Representative images showing switchable electrical coupling between HeLa cells enabled by CarGAP.

To examine the integrated use of CarGAP and NAVI-Cy5 in electrophysiology, we employed HeLa cells, which are substantially larger than N2A cells and thus more amenable to patch clamping. WT HeLa cells possess basal-level gap junctions composed of connexin45 (Cx45)^60^. As a result, the WT HeLa cells surrounding the patched exhibited an average voltage response of 35.9 ± 4.8% relative to the patched cell, suggesting a weak but non-negligible electrical coupling (Fig. 3b,c). Therefore, we created a *Cx45*-knockout cell line, CarGAP HeLa, that stably produces CarGAP. The engineered cells adjacent to the patched cells exhibited an average voltage response of 61.9 ± 7.6 %, substantially higher than that of WT HeLa cells. This finding indicated that CarGAP channels enabled stronger electrical coupling than the native Cx45 channels in HeLa cells.

The voltage response of the CarGAP HeLa cells diminished to 36.1 ± 5.7 %, after treatment with 500 μM AdoB_12_ for 6 hours, in contrast, the response remained largely unchanged (58.4 ± 7.0 %) when cultured with 500 μM MeB_12_, demonstrating that AdoB_12_-induced CarH_C_ oligomerization is essential for blocking CarGAP channels and decoupling the electrical signals among the cells (Fig. 3b,c). Upon green light exposure (3 W/cm^2^, 5 min), the electrical coupling was restored to the level observed in the absence of AdoB_12_, with a comparable voltage response of 58.3 ± 7.5% (Fig. 3b,c). These results showcased the reversible control and facile optical recording of dynamic electrical coupling among interconnected cells enabled by the combination of the chemo-optogenetic tool CarGAP and the far-red hybrid voltage indicator NAVI-Cy5, pointing a novel approach to studying electrophysiology in complex multicellular structures.

WT HeLa refers to wild-type HeLa cells, while CarGAP HeLa indicates Cx45 knockout HeLa cells stably expressed with Cx43-CarH_C_. + MeB_12_ or + AdoB_12_ refers to cells treated with 500 μM MeB_12_ or AdoB_12_ for 6 hours. + AdoB_12_ + 570 nm refers to those treated with AdoB_12_ and then exposed to 5 minutes of illumination from a Xenon lamp equipped with a 550-590 nm filter (∼3 W/cm²). Patched cells, indicated by red dashed lines, were electrically stimulated with a triangle waveform ranging from -100 to +50 mV at a frequency of 10 Hz. Corresponding Cy5 fluorescence intensities (excitation at 637 nm, 1 W/cm²) were normalized from 0 (-100 mV) to 100% (+50 mV). Scale bar: 20 µm. **c**, Quantification of electrical coupling between patched cells and adjacent cells under various conditions. Data are presented as mean ± SEM (n = 30, 25, 23, 31, and 28 adjacent cells). A two-sided t-test was used. * p < 0.05; ** p < 0.01; n.s., not significant.

### Chemo-optogenetically controlled intercellular cAMP transport in the *Drosophila* germarium

While CarGAP has shown its might in controlling mass transfer and electrical coupling among adjacent mammalian cells in vitro, it remains to be seen whether this tool can also exert a delicate control over cellular signaling in more complex biological systems. Gap junctions in invertebrates such as *Drosophila* mainly consist of innexins, of which two octameric complexes docked together to form a gap junction channel^61^. In the *Drosophila* ovary, inner germarial sheath (IGS) cells form the niche for controlling germline stem cell (GSC) self-renewal and differentiation^62^; heterotypic gap junctions composed of niche-specific Inx2 and germ cell-specific Inx4 transport cyclic AMP (cAMP) from niche cells to GSCs and their progeny, thereby orchestrating stepwise oocyte differentiation and development^5^. We envisioned that the ability to dynamically control the flow of crucial second messengers such as cAMP via gap junctions in vivo will open a new synthetic biology approach for biological regulation. To explore this possibility, we constructed two CarGAP systems, i.e., niche-specific Inx2-CarGAP (Fig. 4a) and GSC/progeny-specific Inx4-CarGAP (Fig. 5a), by genetically fusing CarH_C_ to the C-termini of Inx2 and Inx4, respectively. To functionally replace the native IGS-specific Inx2 with Inx2-CarGAP in the *Drosophila* ovary, we created transgenic flies harboring the RNAi-resistant (RR) gene construct, *UASz-inx2^RR^-FLAG-carH_C_* (Fig. 4b).

**Fig. 4.**
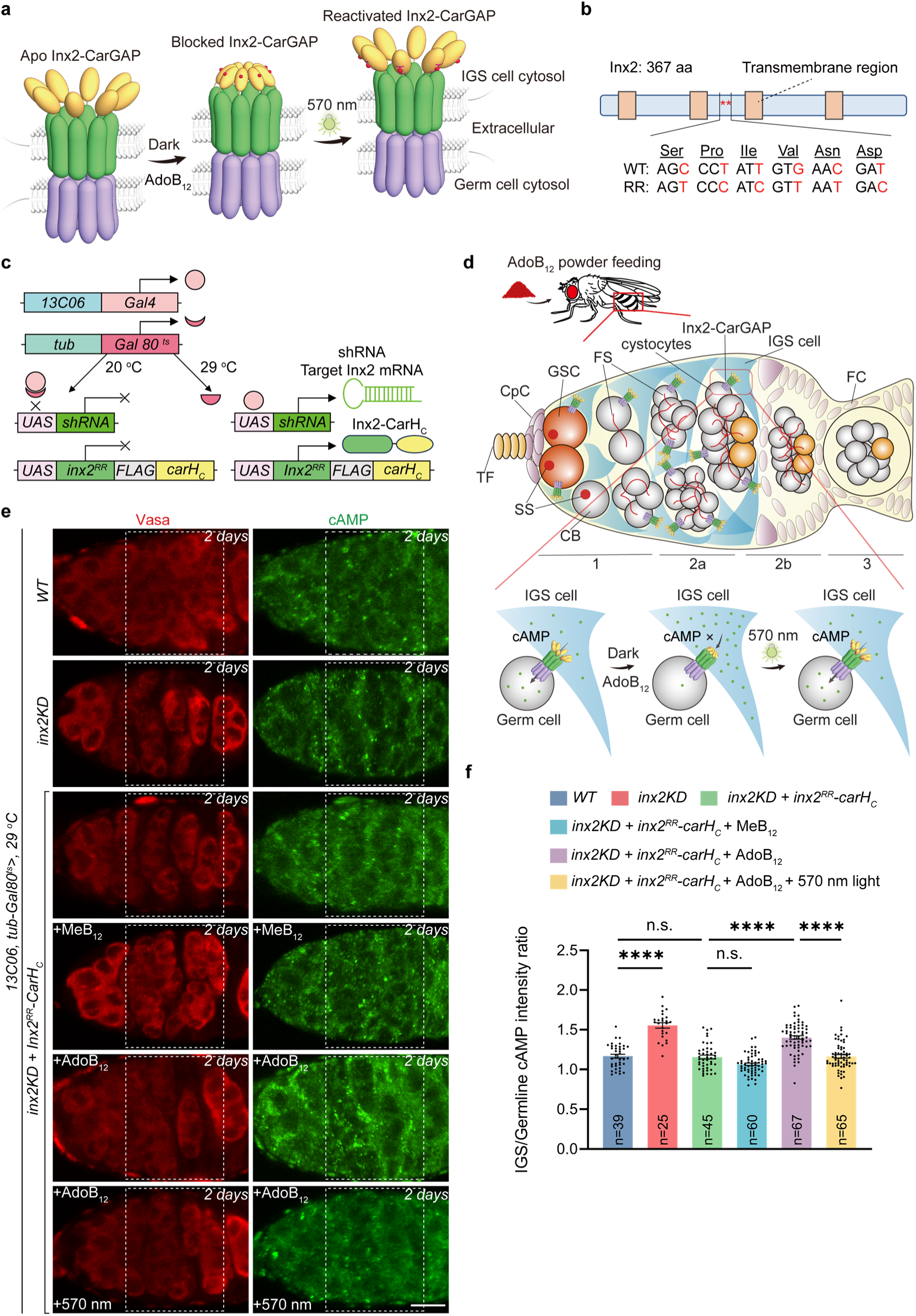
Chemo-optogenetically controlled intercellular transport of cAMP in *Drosophila* germaria enabled by Inx2-CarGAP. **a**, Schematic representation of Inx2-CarGAP. Eight CarH_C_ monomers assemble into two tetramers upon binding to AdoB_12_, blocking the heterotypic gap junction channel within the IGS cell cytosol, while green light (570 nm) reactivates the channel. **b**, Domains of the Inx2 protein and the design of the RNAi-resistant *inx2* gene replacement. Codons in the shRNA target region (BDSC #80409) are replaced with synonymous codons. **c**, Schematic representation of *13C06-Gal4* and *tub-Gal80 ^ts^* for expressing Inx2-CarGAP in *Drosophila* ovaries. The expressions of *Gal4* and *Gal80^ts^*are driven by *13C06^ts^*, a tissue-specific promoter strongly expressed in IGS cells, and *tub*, a constitutively active tubulin promoter, respectively. Gal80^ts^ is a temperature-sensitive inhibitor of the transcriptional activator Gal4. At low temperatures (e.g., 20°C), Gal80^ts^ binds to Gal4, suppressing the transcription of *inx2 shRNA* and *inx2-carH_C_*. At elevated temperatures (e.g., 29°C), Gal80^ts^ dissociates from Gal4, initiating the transcription of *UAS-inx2RNAi* and *UASz-inx2^RR^-carH_C_*. **d**, Schematic representation of Inx2-CarGAP-controlled cAMP transport from IGS cells to germ cells within a *Drosophila* germarium. Under dark conditions, the transgenic fruit fly is fed AdoB_12_ powders, which induce blockage that prevents cAMP from exiting IGS cells. Green light illumination reopens the gap junction channels on the side of IGS cells, restoring cAMP transport between IGS cells and germline. **e**, Single-section confocal images showing Inx2-CarGAP-controlled cAMP transport from IGS cells to germ cells. cAMP is immunostained with a rabbit monoclonal anti-cAMP antibody (green channel), and the germline is stained with an anti-vasa antibody (red channel). Compared to *13C06^ts^*-driven *WT* controls, cAMP levels decreased in germ cells in the *inx2KD* and *inx2KD+inx2-carH_C_*+AdoB_12_ (in the dark) groups. However, cAMP levels were restored in *inx2KD+inx2-carH_C_*, *inx2KD+inx2-carH_C_*+MeB_12_ (in the dark), and *inx2KD+inx2-carH_C_*+AdoB_12_ followed by 5 minutes of green light irradiation (80 mW/cm^2^). All groups were cultured at 29°C to induce the production of Inx2-CarGAP. Scale bar: 10 µm. **f**, Quantification of relative cAMP levels (IGS vs. germline) in **e**. Detailed quantification description was shown in Supplementary Fig. 8. Data are presented as mean ± SEM (n = number of germaria). Statistical analysis was performed using repeated-measures one-way ANOVA with Dunnett’s test. ****p ≤ 0.0001; n.s., not significant.

**Fig. 5.**
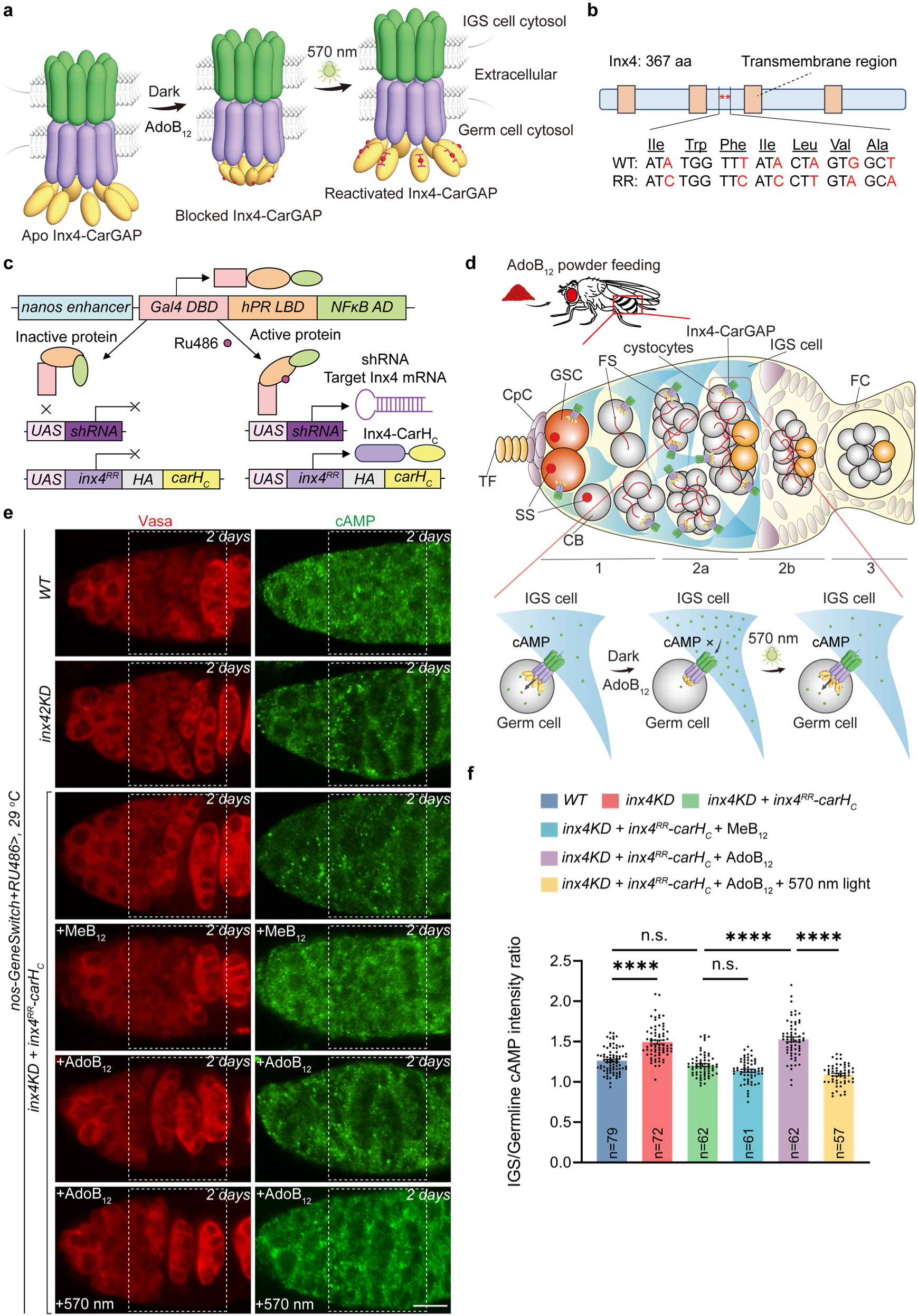
Chemo-optogenetically controlled intercellular transport of cAMP in *Drosophila* germaria enabled by Inx4-CarGAP. **a**, Schematic representation of Inx4-CarGAP. Eight CarH_C_ monomers assemble into two tetramers upon binding to AdoB_12_, blocking the heterotypic gap junction channel within the germ cell cytosol, while green light (570 nm) reactivates the channel. **b**, Domains of the Inx4 protein and the design of the RNAi-resistant *inx4* gene replacement. Codons in the shRNA target region (BDSC #77147) are replaced with synonymous codons. **c**, Schematic representation of *nos-GeneSwitch-Gal4* for expressing Inx4-CarGAP in *Drosophila* ovaries. The expression of *GeneSwitch-Gal4* is driven by the germline-specific nanos promoter. The GeneSwitch-Gal4 protein activates the expression of *UAS-inx4RNAi* and *UASz-inx4^RR^- carH_C_* only in the presence of RU486. **d**, Schematic representation of Inx4-CarGAP-controlled cAMP transport from IGS cells to germ cells within a *Drosophila* germarium. Under dark conditions, the transgenic fruit fly is fed AdoB_12_ powders, which induce blockage that prevents cAMP from entering germ cells. Green light illumination reopens the gap junction channels on the side of germ cells, restoring cAMP transport between IGS cells and germline. **e**, Single-section confocal images showing Inx4-CarGAP-controlled cAMP transport from IGS cells to germ cells. cAMP is immunostained with a rabbit monoclonal anti-cAMP antibody (green channel), and the germline is stained with an anti-vasa antibody (red channel). Compared to those in *nos-GeneSwitch-Gal4*-driven wild-type (WT) controls, cAMP levels decreased in germ cells in the *inx4KD* and *inx4KD+inx4-carH_C_+*AdoB_12_ (in the dark) groups. However, cAMP levels were restored in *inx4KD+inx4-carH_C_*, *inx4KD+inx4-carH_C_+*MeB_12_, and *inx4KD+inx4-carH_C_+*AdoB_12_ followed by 5 minutes of green light irradiation (80 mW/cm²). All groups were cultured at 29 °C, consistent with the control culture conditions used in the Inx2-CarGAP experiments. Scale bar: 10 µm. **f**, Quantification of relative cAMP levels (IGS cells vs. germline) in **e**. Detailed quantification description was shown in Supplementary Fig. 8. Data are presented as mean ± SEM (n = number of germaria). Statistical analysis was performed using repeated-measures one-way ANOVA with Dunnett’s test. ****p ≤ 0.0001; n.s., not significant.

To verify the expression of Inx2-FLAG-CarH_C_ in IGS cells, we first employed the somatic cell-specific c587-Gal4 driver (Supplementary Fig. 6a)^63^. For enhanced experimental flexibility in fly crosses, we subsequently developed a thermal induction system using the *13C06-Gal4* driver, which similarly targets somatic cells, including IGS cells, follicle stem cells (FSCs), and prefollicle cells adjacent to the region 2a/2b border^64^. In this system, denoted as *inx2KD+inx2^RR^-carH_C_*, the transcription factor Gal4 is driven by the 13C06 promoter. At an ambient temperature of 20°C, Gal4 expression is repressed by Gal80^ts^. However, at an elevated temperature of 29°C, Gal80^ts^ repression is lifted, allowing Gal4 to activate the expression of *inx2^RR^-FLAG-CarH_C_* through its upstream activating sequence (UAS). Concurrently, *inx2 shRNA* is co-expressed to selectively knock down the native *inx2* gene (Fig. 4c). In a similar vein, to incorporate the GSC/progeny-specific Inx4-CarGAP into the *Drosophila* ovaries, we first use the germline specific *nos-Gal4* driver^65^ to verify the expression of Inx4-HA-CarH_C_ in germ cells. Further, we made the transgenic flies to possess the gene constructs, including *UASz-inx4^RR^-carH_C_* and *UAS-shRNA*, along with the germline-specific *nos-GeneSwitch-Gal4* driver (mainly express in 2a region during germarium development) to achieve the genetic replacement of the endogenous *inx4* to *inx4-carH_C_* (Fig. 5b,c and Supplementary Fig. 6b). The GeneSwitch-Gal4, which combines the Gal4/UAS system with a receptor of RU486 (mifepristone)^66^, allowed us to activate the expression of *inx4^RR^-carH_C_* in response to the administration of RU486, while co-expressing *inx4 shRNA* to silence the native *inx4* gene (Fig. 5c).

To assess the influence of CarGAP on relative distributions of cAMP under various conditions, we used a commercial rabbit anti-cAMP monoclonal antibody to fluorescently stain and quantify this second messenger in the ovaries. The relative fluorescence intensities of cystocyte clusters and their surrounding IGS cells would reflect the transferability of cAMP from IGS cells (somatic cells) to the germline (GSCs, cystoblasts [CBs], cystocytes) (Fig. 4d,5d). Detailed methods for quantifying cAMP fluorescence intensity in IGS/germline cells are provided in Supplementary Fig. 8. Consistent with the previous finding that gap junctions are essential for the cAMP transport in the ovaries^5^, knocking down either *inx2* or *inx4* disrupted the cAMP transport from IGS cells to germ cells, as evidenced by the contrast in fluorescence between these cells (Fig. 4e,5e). Expression of Inx2-CarGAP in the *inx2KD* germarium (i.e., *inx2KD+inx2^RR^-carH_C_*) or Inx4-CarGAP in the *inx4KD* germarium (i.e., *inx4KD+inx4^RR^-carH_C_*) restored the flow of cAMP from IGS cells to germ cells, confirming the functionality of the Inx-CarGAP channels (Fig. 4d-f,5d-f). Given the bidirectional nature of gap junctions, IGS-specific Inx2-CarGAP and Germline-specific Inx4-CarGAP should be functionally indistinguishable and chemo-optogenetic control over the cAMP transport may be accomplished from either side, IGS cells or Germline (Fig. 4d,5d). Consistent with this prediction, orally feeding both transgenic flies (i.e., *inx2KD+inx2^RR^-carH_C_* and *inx4KD+inx4^RR^-carH_C_*) with AdoB_12_ for two days stemmed the flow of cAMP from IGS cells to germ cells, leading to its accumulation and depletion in IGS cells and germ cells, respectively. By contrast, feeding the flies with MeB_12_ failed to do so, suggesting that AdoB_12_-induced CarH_C_ oligomerization is essential for the observed inhibition of CarGAP channels in vivo (Fig. 4d-f,5d-f). To examine whether light-induced protein disassembly can restore the cAMP transport, we placed these transgenic flies under green light irradiation (570 nm, 80 mW/cm^2^) (Supplementary Fig. 7). It turned out that brief light exposure for merely 5 min was sufficient to increase cAMP in germ cells to a level comparable to that of the control, strongly suggesting the resumed flow of cAMP among the cells (Fig. 4d-f,5d-f). These results highlighted CarGAP as a robust chemo-optogenetic tool for reversibly controlling the intercellular transport of the crucial second messenger cAMP in complex biological systems.

### Modulating ovarian development via CarGAP

Tu and coworkers recently showed that IGS-specific Inx2 proteins form gap junctions with GSC/progeny-specific Inx4 proteins in the *Drosophila* ovary, which further dictate stepwise GSC lineage development, including GSC self-renewal, germline cyst formation, meiotic double-strand DNA break (DSB) formation, and oocyte specification^5^. While *inx2* knockdown (i.e., *inx2KD*) led to pronounced phenotypic defects in the ovaries, such as fewer GSCs, more CBs, decreased *H2AvD* expression, and double oocytes in stage 1 egg chambers, the expression of *inx2^RR^- FLAG-carH_C_* in the *inx2KD* background (i.e., *inx2KD+inx2^RR^-carH_C_*) at 29°C restored these ovarian phenotypes to a level comparable to the WT control, supporting the normal function of Inx2-CarGAP in vivo (Fig. 6a-d). Interestingly, simply feeding the *inx2KD+inx2^RR^-carH_C_* fruit flies with AdoB_12_ powders, but not MeB_12_, sufficiently caused severe ovarian defects in GSC maintenance, CB differentiation, meiotic DSB formation and oocyte determination akin to *inx2KD*, showing that AdoB_12_-induced CarH_C_ oligomerization can efficiently block the gap junction channels in vivo (Fig. 6a-d). Reminiscent of the Inx2-CarGAP system, the production of Inx4-CarH_C_ in *inx4KD+inx4^RR^-carH_C_* proved to be able to rescue the defective development of GSCs and their progeny caused by *inx4KD*, indicative of functional Inx4-CarGAP channels formed by Inx4-CarH_C_ and Inx2 (Fig. 6e-h). Feeding the *inx4KD+inx4^RR^-carH_C_* flies with AdoB_12_, but not MeB_12_, induced ovarian defects, such as GSC loss, CB accumulation, defective meiotic DSB formation and oocyte determination, comparable to *inx4* knockdown (i.e., *inx4KD*) (Fig. 6e-h). It is noteworthy that the developmental anomalies caused by AdoB_12_-induced CarGAP blockage were somewhat milder than those by gene silencing in *inx2KD* and *inx4KD,* the latter of which exhibited severe aberrations in germarium shape, including drastic reduction in size and loss of contour (Fig 6a,e). The phenotypic differences between chemogenetic inhibition and gene knockout might stem from two possible mechanisms. First, the AdoB_12_-induced gap junction blockage may incompletely suppress metabolite transfer through innexin channels due to their larger pore size compared to connexins. Second, it is very likely that the pleiotropic functions of innexins beyond channel activity, including those in cell tiling^67^, cell adhesion^68^, and cytoskeletal stabilization^69^, still persist even when channel function is chemically inhibited, which may contribute to distinct phenotypic outcomes^70^. These results established CarGAP, with oral administration of AdoB_12_, as a convenient molecular tool for controlling gap junctions in vivo.

**Fig. 6.**
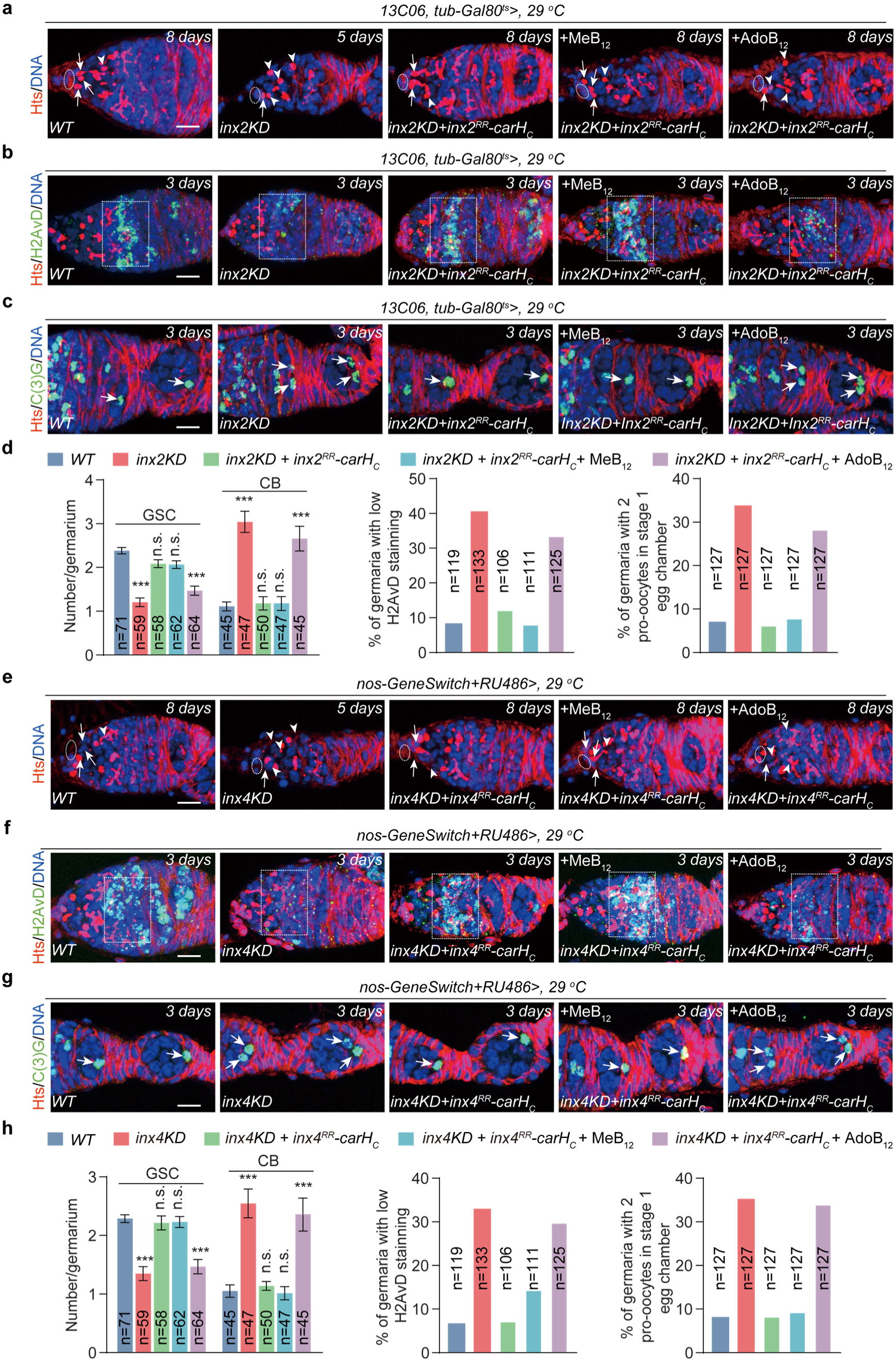
AdoB_12_-induced gap junction blockage in *Drosophila* germaria. **a**, GSC and CB populations in *Drosophila* ovaries under various conditions. Hts is a marker for the spherical spectrosome in GSCs and CBs, as well as the branched fusome in cysts. Cap cells are delineated by dashed lines, and germ cells in proximity to the cap cells are classified as GSCs. Compared to the control (*13C06^ts^-driven WT*), the numbers of GSCs (indicated by arrows) decreased, while the numbers of CBs (indicated by arrowheads) increased in both *inx2KD* and *inx2KD+inx2-carH_C_*+AdoB_12_, but were restored in both *inx2KD+inx2-carH_C_* and *inx2KD+inx2-carH_C_*+MeB_12_. All groups were cultured at 29 °C to induce the production of Inx2-CarGAP. Scale bar: 10 µm. **b**, Expression of H2AvD in meiotic 16-cell cysts. Compared to the control (*13C06^ts^*-driven *WT*), H2AvD expression decreased in *inx2KD* and *inx2KD+inx2-carH_C_*+AdoB_12_, but was restored in both *inx2KD+inx2-carH_C_* and *inx2KD+inx2-carH_C_*+MeB_12_. All groups were cultured at 29 °C for induction. Scale bar: 10 µm. **c**, Incidence of oocytes in stage 1 egg chambers. Compared to the control (*13C06^ts^*-driven *WT*), the incidence of oocytes increased in *inx2KD* and *inx2KD+inx2-carH_C_*+AdoB_12_, and was restored in *inx2KD+inx2-carH_C_* and *inx2KD+inx2-carH_C_*+MeB_12_. All groups were cultured at 29 °C for induction. Scale bar: 10 µm. **d**, Quantification results for **a** (left), **b** (middle), and **c** (right). Data are presented as mean ± SEM (n = number of germaria). Statistical analysis was performed using Student’s t-test (***p ≤ 0.001; n.s., not significant). **e**, GSC and CB populations in *Drosophila* ovaries under various conditions. Hts is a marker for the spherical spectrosome in GSCs and CBs, as well as the branched fusome in cysts. Cap cells are delineated by dashed lines, and germ cells in proximity to the cap cells are classified as GSCs. Compared to the control (WT), the numbers of GSCs (indicated by arrows) decreased, while the numbers of CBs (indicated by arrowheads) increased in *nos-GeneSwitch-Gal4*-driven *inx4KD* and *inx4KD+inx4-carH_C_*+AdoB_12_, but were restored in both *inx4KD+inx4-carH_C_* and *inx4KD+inx4-carH_C_*+MeB_12_. All groups were cultured at 29 °C, consistent with those used in the Inx2-CarGAP experiments. Scale bar: 10 µm. **f**, Expression of H2AvD in meiotic 16-cell cysts. Compared to the control (*nos-GeneSwitch-Gal4*-driven *WT*), H2AvD expression decreased in *inx4KD* and *inx4KD*+*inx4-carH_C_*+AdoB_12_, but was restored in both *inx4KD+inx4-carH_C_* and *inx4KD+inx4-carH_C_*+MeB_12_. All groups were cultured at 29 °C, consistent with those used in the Inx2-CarGAP experiments. Scale bar: 10 µm. **g**, Incidence of oocytes in stage 1 egg chambers. Compared to the control (*nos-GeneSwitch-Gal4*-driven *WT*), the incidence of oocytes increased in *inx4KD* and *inx4KD*+*inx4-carH_C_* +AdoB_12_, and was restored in *inx4KD+inx4-carH_C_* and *inx4KD+inx4-carH_C_*+MeB_12_. All groups were cultured at 29 °C, consistent with those used in the Inx2-CarGAP experiments. Scale bar: 10 µm. **h**, Quantification results for **e** (left), **f** (middle), and **g** (right). Data are presented as mean ± SEM (n = number of germaria). Statistical analysis was performed using Student’s t-test (***p ≤ 0.001; n.s., not significant).

## Discussion and Conclusions

Since the first electron micrograph of gap junctions was obtained over sixty years ago^71^, researchers have sought to understand their critical roles in rapid signal transduction and electrical coupling across multicellular structures. However, reliable tools for noninvasive control and interrogation of these intercellular channels remain elusive. While optogenetic approaches offer theoretical precision, practical limitations persist: light delivery challenges in vivo, and problematic dark-state activity in common tools like rhodopsins and Cry2 domains that often compromises data accuracy^72^. Chemogenetic methods using small molecules can circumvent some delivery issues, but existing standards like rapamycin present their own constraints. Though widely adopted for FKBP-FRB dimerization *in vitro*^73^ and *in vivo*^74^, rapamycin’s inhibition of mTOR signaling^75, 76^—which directly impacts the trafficking of gap junction proteins^77^—limits its utility for studying intercellular communication.

CarGAP combines the strengths of both approaches. Built around the compact (∼23 kDa; comparable to GFP), B_12_-dependent photoreceptor CarH_C_, this system leverages the unique advantages of vitamin B_12_: (1) marked green-light sensitivity (10 mW/cm²— at least three orders of magnitude lower than conventional CALI methods^29, 30^) (2) exceptional biocompatibility^54^, and (3) the ability to cross the blood-brain barrier when administered orally^78^. Unlike signaling-active molecules, vitamin B_12_ has not been known for any pronounced role in biological signaling and mainly serves as a nutrient, particularly crucial for neural function^78^. By integrating the precision of optogenetics with the tissue penetration of chemogenetics, CarGAP may finally enable rigorous investigation of GJC-mediated signaling in complex biological systems, including the nervous system.

This study has demonstrated the B_12_-dependent CarGAP system as a versatile chemo-optogenetic tool for investigating GJC-mediated signaling in mammalian cell cultures and *Drosophila* ovaries. In N2A cells, connexin-based CarGAP enabled precise control of intercellular 2’,3’-cGAMP transport and subsequent cGAS-STING activation in neighboring cells, a crucial process in bystander immunity. When combined with the far-red fluorescent voltage indicator NAVI-Cy5, this system permitted rapid, reversible modulation and optical monitoring of electrical coupling in cell clusters. The conserved architecture of gap junction proteins suggests broad applicability of the CarGAP design principle across different biological systems.

We successfully adapted CarGAP to invertebrate gap junctions, creating an innexin-based version that regulates cAMP flow from somatic niche cells to germline in *Drosophila* ovaries. This enabled stepwise control of germ cell development. Unlike genetic approaches that eliminate all protein functions, AdoB_12_-induced CarH_C_ oligomerization specifically blocks CarGAP’s channel activity while preserving non-channel roles (e.g., structural scaffolding and cell adhesion). This selectivity revealed previously obscured nuances in gap junction function within complex biological systems.

Given the established importance of gap junction-mediated signaling in mammalian development and disease^79, 80^, CarGAP represents a powerful synthetic biology tool. Its vitamin B_12_ dependence provides a simple, unified control mechanism for studying complex developmental processes and disease mechanisms, eliminating the need for a myriad of induction methods. This system thus opens new avenues for investigating intercellular communication across diverse biological contexts.

In summary, we have developed CarGAP, a versatile molecular tool that enables reversible control of GJCs across biological systems - from mammalian cell cultures to *Drosophila* ovaries in vivo. This system centers on the chemo-optogenetic switch CarH_C_, which undergoes AdoB_12_-dependent oligomerization and light-triggered disassembly. By combining vitamin B_12_ administration with green light illumination, CarGAP effectively modulates both chemical signal (including second messengers like 2’,3’-cGAMP and cAMP) and electrical coupling between connected cells. Our application of this tool has revealed fundamental aspects of heterotypic gap junction function during *Drosophila* ovarian development. The generalizable design principle of CarGAP promises to advance our understanding of intercellular communication mechanisms that are fundamental to multicellular life.

## Material and Methods

### Plasmid construction

All plasmids used in this study are summarized in Supplementary Table 1. The His-tagged *carH_C_* gene was initially inserted into the pET-22b(+) vector for protein expression in *E. coli*. The gene encoding *Cx43-carH_C_* was codon-optimized for the mammalian cell expression, obtained through chemical synthesis (GENEWIZ), and inserted between KpnI and BamHI sites in the pcDNA3.1(+) vector (Invitrogen). To construct the fly transgenic expression vector, the Q5® Site-Directed Mutagenesis Kit (NEB) was used to generate *UASz-inx2^RR^-FLAG* and *UASz-inx4^RR^-HA*. The *carH_C_* gene was inserted into *UASz-inx2^RR^-FLAG* and *UASz-inx4^RR^-HA* using Gibson assembly with NEBuilder® HiFi DNA Assembly Master Mix (NEB). All relevant genetic components were confirmed by sequencing (BGI Genomics Co., Ltd.). *Escherichia coli* strain DH5α was used for plasmid amplification, followed by plasmid extraction using the SanPrep Endotoxin-Free Plasmid Mini Kit (Sangon).

### Protein expression

Protein expression was conducted using *E. coli* strain BL21(DE3) as the host. Bacteria were cultured in LB medium at 37°C until reaching mid-log phase (OD600 ∼0.6-0.8). Protein expression was induced with 300 μM isopropyl-β-d-1-thiogalactopyranoside (Sangon Biotech) and continued at 37°C for 4 hours. Bacteria were harvested, and proteins were purified using HisTrap columns (GE Healthcare Inc.). The purified proteins were dialyzed against 5 liters of Milli-Q water at 4°C, repeated six times, and lyophilized at −100°C. The lyophilized proteins were stored at −80°C until use.

### Analytical Size-Exclusion Chromatography (SEC)

Analytical SEC was performed using an ÄKTA Pure system (GE Healthcare) equipped with a Superose 6 Increase 10/300 GL column. Protein samples (30 μM), with or without 120 μM vitamin B_12_ (AdoB_12_ or MeB_12_), were loaded onto the column pre-equilibrated with PBS.

### Cell culture and transfection

N2A cells (CCL-131), HEK293T (CRL-3216), and HeLa cells (CCL-2) were obtained from ATCC. Cells were cultured in DMEM (Gibco) supplemented with 0.5% penicillin-streptomycin (Gibco) and 10% FBS (Gibco) in a humidified incubator with 5% CO_2_ at 37 °C. Transient transfections were performed using Lipofectamine 3000 reagent (Thermo Fisher) mixed with 2000 ng of the plasmid per 35 mm cell culture dish (SPL Life Sciences) for 36 hours. HT-DNA was transfected at a concentration of 5 μg/ml (D6898, Sigma) for cGAS activation.

### Generation of stable cell lines

To generate the Cx43-CarH_C_ N2A stable cell line, we first cloned *Cx43-carH_C_* into the PLVX-PURO vector and then packaged the plasmid into lentiviral particles. N2A cells were infected with the packaged lentivirus, and a pool of positively transduced cells was obtained through puromycin selection. Subsequently, single-cell clones were isolated using flow cytometry, and the mRNA transcription levels of these individual clones were evaluated using qPCR. The N2A Cx43-CarH_C_ stable cell line is maintained in culture medium containing 2 μg/ml puromycin.

To generate the Cx45KO + Cx43-CarH_C_ HeLa stable cell line, synthetic *GJC1 (Cx45) gRNA2* was first cloned into the lentiCRISPR v2 vector between two BsmBI restriction sites^60^. HeLa cells were infected with the packaged lentivirus and selected for positive cell pools using puromycin. Based on the Cx45 KO HeLa stable cell line, blasticidin (BSD) was cloned into the PLVX-PURO-*Cx43-carH_C_* plasmid to replace puromycin, resulting in a BSD-resistant lentiviral vector. The BSD-*Cx43-carH_C_* lentivirus was then used to infect Cx45 KO HeLa cell lines, and positive cell pools were obtained through BSD resistance selection. Single-cell clones were isolated using flow cytometry, and the mRNA transcription levels of these individual clones were evaluated using qPCR. The Cx45KO + Cx43-CarH_C_ stable cell line is maintained in culture medium containing 2 μg/ml puromycin and 2 μg/ml BSD.

### Dye transfer assays

N2A cells were trypsinized in TrypLE (Gibco) for 3 minutes, suspended in culture medium, centrifuged at 200 g for 5 minutes, and then resuspended in 1.5 ml of culture medium containing a staining solution with 5 μM DiD (Dil) and 5 μM calcein AM. The cells were incubated at 37 °C for 20 minutes. After incubation, the cells were washed with phosphate-buffered saline (PBS), centrifuged three times, and co-cultured with unstained recipient cells at a ratio of 1:40 (donor cells: recipient cells). Cells required at least 5 hours to fully attach to the confocal dishes for high-quality imaging. Fluorescent imaging was then performed using an Inverted Microscope Ti2-E (Nikon) equipped with an X-Cite 110 LED Microscope Light Source (Lumen Dynamics). The 488 nm laser (10 mW) was set to 5% intensity, and emission was collected at 515–555 nm for the green channel; the 633 nm laser (10 mW) was set to 20% intensity, and emission was collected at 663–738 nm for the red channel. All vitamin B_12_-related cell culturing was conducted under dim red LED light (660 nm) to prevent photocleavage.

### FACS analyses

After co-culturing for 5 or 10 hours (donor cells: recipient cells = 1:20 or 1:40), the cells were detached and resuspended in PBS. To assess the dye diffusion ratio for GarGAP, cells were analyzed using 488 nm, 561 nm and 633 nm lasers equipped with 530/30 (for calcein), 582/15 (for Dil) and 660/20 (for DiD) emission filters. A total of 10,000 or 20,000 single cells were recorded to analyze the diffusion ratio to screen the optimal co-culture time. A total of 20,000 single cells were recorded to analyze the diffusion ratio to assess CarGAP functionality. Data analysis was performed using FlowJo X (version 10.8.1).

### Digitonin permeabilization

Cells were treated with 2 µg/ml of 2′3′-cGAMP in 1 ml of digitonin solution [50 mM HEPES, pH 7.0; 100 mM KCl; 3 mM MgCl_2_; 0.1 mM dithiothreitol (DTT); 85 mM sucrose; 0.2% BSA; 1 mM ATP; 0.1 mM GTP; 1 µg/ml digitonin] at 37 °C for 30 minutes. After treatment, the cells were incubated in fresh DMEM for imaging.

### Control and fluorescent imaging of electrical coupling

To control and monitor electrical coupling with CarGAP and NAVI-Cy5, the Cx43KO + Cx43-CarH_C_ HeLa stable cell line was seeded onto 12-mm glass coverslips coated with Matrigel® Matrix and transfected with NAVI at approximately 80% confluency. To chemically block Cx43-CarH_C_, the cells were treated with 500 μM vitamin B_12_ (AdoB_12_ or MeB_12_) dissolved in DMEM with 10% FBS, away from light, for about 6 hours. To create the cell membrane-bound voltage indicator, NAVI-Cy5, 100 nM Cy5-conjugated nanobody was added to the cell culture medium 2 hours before voltage imaging. Subsequently, the cells were rinsed with Tyrode’s buffer (imaging solution) to remove excess Cy5-nanobody. As to green light illumination, cells were illuminated with a Xenon lamp equipped with a 550-590 nm filter at approximately 3 W/cm² for 5 minutes.

For single-cell patch clamp, borosilicate glass electrodes (Sutter) were pulled to a tip resistance of 2.5–5 MΩ and filled with an internal solution (pH 7.3 and 295 mOsm/kg) containing 125 mM potassium gluconate, 8 mM NaCl, 0.6 mM MgCl_2_, 0.1 mM CaCl_2_, 1 mM EGTA, 10 mM HEPES, 4 mM Mg-ATP, and 0.4 mM GTP·Na_2_. The cells were clamped using an Axopatch 200B (Axon Instruments) amplifier at -30 mV at room temperature. The patched cells were subjected to a triangle waveform from -100 mV to +50 mV for 6 seconds. Voltage imaging was performed simultaneously at 10 Hz with 4×4 pixel binning under illumination from a 637 nm laser beam at ∼ 1 W/cm². The imaging experiments were conducted with an inverted fluorescence microscope (Nikon TiE), equipped with a 40× 1.3 NA oil immersion objective lens, a 637 nm laser line (Coherent OBIS), and a scientific CMOS camera (Hamamatsu ORCA-Flash 4.0 v2), and the LabVIEW software (National Instruments, version 15.0). Images were captured at 2×2 pixel binning with a 100-ms exposure time and analyzed using ImageJ/Fiji (version 1.52d).

### Construction of transgenic *Drosophila* strains

The *attB* site sequence (5’-GGGTGCCAGGGCGTGCCCTTGGGCTCCCCGGGCGCGT-3’) is present in the *UASz-inx2^RR^-carH_C_* and *UASz-inx4^RR^-carH_C_* plasmids. The DNA constructs were integrated into the *attP40* site on the second chromosome or into the *attP2* site on the third chromosome via PhiC31-mediated transgenesis.

### *Drosophila* culture

The *Drosophila* stocks used in this study are described in FlyBase, unless otherwise specified: *13C06-Gal4*, *nos-Gal4*, *tubulin-Gal80^ts^*, *inx2* RNAi (BL29306 and BL80409), *inx4* RNAi (BL35607 and BL77147). Flies were maintained and crossed at room temperature on standard cornmeal/molasses/agar media unless otherwise specified. To maximize the effect of *inx2* RNAi-mediated knockdown and *inx2-carH_C_* gene overexpression, newly eclosed flies were shifted to 29 °C for the specified number of days before analyzing ovarian phenotypes. To maximize the effect of *inx4* RNAi-mediated knockdown and *inx4-carH_C_* gene overexpression, newly eclosed flies were fed with Ru486 for the specified number of days before analyzing ovarian phenotypes. Some experiments were conducted under dim red LED light (660 nm) to prevent photocleavage of vitamin B_12_ when necessary.

### AdoB_12_ supplementation and light illumination

In the dye transfer and cGAMP transport assays, prepared donor and recipient cells were mixed and incubated in DMEM culture medium supplemented with 500 μM vitamin B_12_ (either AdoB_12_ or MeB_12_) for 5 hours. For the *Drosophila* vitamin B_12_ experiments, 0.01 g of vitamin B_12_ red powder (AdoB_12_ or MeB_12_) was evenly distributed on the food in each tube (Supplementary Fig. 7a). Flies were cultured in tubes covered with aluminum foil to ensure dark conditions.

Cold LEDs purchased from PURI Materials (Shenzhen) were used as the green light source (570 nm) (excluding electrophysiology experiments). For the dye transfer and cGAMP transport assays in mammalian cells, the cell culture plate was illuminated with green LED light at an intensity of 10 mW/cm^2^ for 5 minutes before fluorescent imaging. For the CarGAP experiment in *Drosophila*, live flies were placed in transparent tubes and exposed to green LED light at an intensity of 80 mW/cm^2^ for 5 minutes (Supplementary Fig. 7b), followed by ovary dissection and fixation.

### Immunohistochemistry and confocal imaging

Ovaries were dissected at room temperature in Grace’s medium, fixed with 4% paraformaldehyde, and then stained according to established protocols^5^. The following antibodies were used: Rabbit anti-pSTING (1:50, Cell Signaling, #62912), rabbit anti-cGAS (1:100, Cell Signaling, #79978), mouse anti-FLAG (1:1000, Sigma-Aldrich, F1804), rabbit anti-HA (1:200, Sigma-Aldrich, H6908), mouse monoclonal anti-Hts (1:50, DSHB, 1B1), rat monoclonal anti-Vasa (1:50, DSHB, anti-vasa), rabbit polyclonal anti-pS137 γ-H2Av (1:1000, Rockland, #600-401-914), rabbit polyclonal anti-C3G (1:10000, gift from Lilly, NICHD/DIR), and rabbit monoclonal anti-cAMP (1:200, Abcam, ab134901). For the rabbit polyclonal anti-Aub antibody, a peptide (MHKSEGDPRGSVRGC) from *Drosophila* Aub was synthesized and injected into a rabbit by GeneScript USA Inc., and the Protein A/G purified IgG was used for staining (1:1000).

### Quantification and statistical analysis

For GSC quantification, spectrosome-containing single germ cells attached to cap cells were counted as GSCs. Statistical analysis was performed using Microsoft Excel and Student’s t-test. For CB quantification, single germ cells with a round fusome that were not attached to cap cells were counted as CBs, with statistical analysis conducted using Student’s t-test in Microsoft Excel. Fluorescence intensity quantification was performed using the embedded software of the Nikon Ti2 confocal microscope. The area of interest was highlighted, and the mean fluorescence intensity ratio between IGS cells and germline (Vasa channel) was exported for all areas of interest (Supplementary Fig. 8). Statistical analysis was conducted using ordinary one-way ANOVA, followed by Tukey’s multiple comparisons test.

## Acknowledgements

This work was supported by the Natural Science Foundation of China Excellent Young Scientists Fund (22122707, F.S.), the Ministry of Science and Technology (2020YFA0908100, F.S.), the Research Institute of Tsinghua, Pearl River Delta (#RITPRD21EG01, F.S.), the Research Grants Council of Hong Kong SAR (GRF Grants #16103519 and #16103421, F.S.), the Research Fellow Scheme (RFS2324-6S05, F.S.), the Young Collaborative Research Fund (C6001-23Y, F.S.), and the Theme-based Research Scheme (T13-602/21-N, T.X.).

## Author contributions

F.S., T.X., P.Z., and R.T. conceived the project and provided supervision. D.C. designed CarGAP. D.C. and M.Z. designed and performed the dye transfer assay. D.C. and X.H. designed and conducted the FACS analysis and cGAMP test assay. D.C. and S.L. designed and performed the patch clamp electrophysiology experiments. D.C., R.T., and X.T. designed and conducted the *Drosophila* experiments with assistance from Z.H. and R.X. F.S., D.C., S.L., and X.H. wrote the manuscript. All authors participated in data analysis and reviewed the final manuscript.

## References

1 Kumar, N. M. & Gilula, N. B. The gap junction communication channel. Cell 84, 381–388 (1996).

2 Skerrett, I. M. & Williams, J. B. A structural and functional comparison of gap junction channels composed of connexins and innexins. Developmental neurobiology 77, 522–547 (2017).

3 Welzel, G. & Schuster, S. Connexins evolved after early chordates lost innexin diversity. Elife 11, e74422 (2022).

4 Verselis, V., White, R., Spray, D. & Bennett, M. Gap junctional conductance and permeability are linearly related. Science 234, 461–464 (1986).

5 Tu, R. et al. Gap junction–transported cAMP from the niche controls stem cell progeny differentiation. Proceedings of the National Academy of Sciences 120, e2304168120 (2023).

6 Ablasser, A. et al. Cell intrinsic immunity spreads to bystander cells via the intercellular transfer of cGAMP. Nature 503, 530–534 (2013).

7 Chen, Q. et al. Carcinoma–astrocyte gap junctions promote brain metastasis by cGAMP transfer. Nature 533, 493–498 (2016).

8 Kizana, E., Cingolani, E. & Marban, E. Non-cell-autonomous effects of vector-expressed regulatory RNAs in mammalian heart cells. Gene therapy 16, 1163–1168 (2009).

9 Vachias, C. et al. Gap junctions allow transfer of metabolites between germ cells and somatic cells to promote germ cell growth in the Drosophila ovary. PLoS Biology 23, e3003045 (2025).

10 Neijssen, J. et al. Cross-presentation by intercellular peptide transfer through gap junctions. Nature 434, 83–88 (2005).

11 Jongsma, H. J. & Wilders, R. Gap junctions in cardiovascular disease. Circulation research 86, 1193–1197 (2000).

12 Bloomfield, S. A. & Völgyi, B. The diverse functional roles and regulation of neuronal gap junctions in the retina. Nature Reviews Neuroscience 10, 495–506 (2009).

13 Almoril-Porras, A. et al. Configuration of electrical synapses filters sensory information to drive behavioral choices. Cell (2024).

14 Zhu, Y., Zhu, H. & Wu, P. Gap junctions in polycystic ovary syndrome: Implications for follicular arrest. Developmental Dynamics (2024).

15 Aasen, T., Mesnil, M., Naus, C. C., Lampe, P. D. & Laird, D. W. Gap junctions and cancer: communicating for 50 years. Nature Reviews Cancer 16, 775–788 (2016).

16 Spray, D., Stern, J., Harris, A. & Bennett, M. Gap junctional conductance: comparison of sensitivities to H and Ca ions. Proceedings of the National Academy of Sciences 79, 441–445 (1982).

17 Peracchia, C. Chemical gating of gap junction channels: roles of calcium, pH and calmodulin. Biochimica et Biophysica Acta (BBA)-Biomembranes 1662, 61–80 (2004).

18 Sáez, J. C., Berthoud, V. M., Branes, M. C., Martínez, A. D. & Beyer, E. C. Plasma membrane channels formed by connexins: their regulation and functions. Physiological reviews 83, 1359–1400 (2003).

19 Caveney, S. The role of gap junctions in development. Annual review of physiology 47, 319–335 (1985).

20 Krüger, O. et al. Defective vascular development in connexin 45-deficient mice. Development 127, 4179–4193 (2000).

21 Gabriel, H.-D. et al. Transplacental uptake of glucose is decreased in embryonic lethal connexin26-deficient mice. The Journal of cell biology 140, 1453–1461 (1998).

22 Pereda, A. E. Electrical synapses and their functional interactions with chemical synapses. Nature Reviews Neuroscience 15, 250–263 (2014).

23 Rost, B. R., Wietek, J., Yizhar, O. & Schmitz, D. Optogenetics at the presynapse. Nature neuroscience 25, 984–998 (2022).

24 Karpova, A. Y., Tervo, D. G., Gray, N. W. & Svoboda, K. Rapid and reversible chemical inactivation of synaptic transmission in genetically targeted neurons. Neuron 48, 727–735 (2005).

25 Ojima, K. et al. Coordination chemogenetics for activation of GPCR-type glutamate receptors in brain tissue. Nature communications 13, 3167 (2022).

26 Rozental, R., Srinivas, M. & Spray, D. C. How to close a gap junction channel: Efficacies and potencies of uncoupling agents. Connexin methods and protocols, 447–476 (2001).

27 Juszczak, G. R. & Swiergiel, A. H. Properties of gap junction blockers and their behavioural, cognitive and electrophysiological effects: animal and human studies. Progress in neuro-psychopharmacology and biological psychiatry 33, 181–198 (2009).

28 Desplantez, T., Verma, V., Leybaert, L., Evans, W. & Weingart, R. Gap26, a connexin mimetic peptide, inhibits currents carried by connexin43 hemichannels and gap junction channels. Pharmacological research 65, 546–552 (2012).

29 Tanabe, T. et al. Multiphoton excitation–evoked chromophore-assisted laser inactivation using green fluorescent protein. Nature methods 2, 503–505 (2005).

30 Tour, O., Meijer, R. M., Zacharias, D. A., Adams, S. R. & Tsien, R. Y. Genetically targeted chromophore-assisted light inactivation. Nature biotechnology 21, 1505–1508 (2003).

31 Bulina, M. E. et al. A genetically encoded photosensitizer. Nature biotechnology 24, 95–99 (2006).

32 Zhang, W. et al. Optogenetic control with a photocleavable protein, PhoCl. Nature methods 14, 391–394 (2017).

33 Wang, R., Yang, Z., Luo, J., Hsing, I.-M. & Sun, F. B12-dependent photoresponsive protein hydrogels for controlled stem cell/protein release. Proceedings of the National Academy of Sciences 114, 5912–5917 (2017).

34 Jiang, B. et al. Injectable, photoresponsive hydrogels for delivering neuroprotective proteins enabled by metal-directed protein assembly. Science Advances 6, eabc4824 (2020).

35 Kainrath, S., Stadler, M., Reichhart, E., Distel, M. & Janovjak, H. Green-light-induced inactivation of receptor signaling using cobalamin-binding domains. Angewandte Chemie International Edition 56, 4608–4611 (2017).

36 Mansouri, M. et al. Smart-watch-programmed green-light-operated percutaneous control of therapeutic transgenes. Nature Communications 12, 3388 (2021).

37 Kutta, R. J. et al. The photochemical mechanism of a B12-dependent photoreceptor protein. Nature communications 6, 7907 (2015).

38 Jost, M. et al. Structural basis for gene regulation by a B12-dependent photoreceptor. Nature 526, 536–541 (2015).

39 Padmanabhan, S., Jost, M., Drennan, C. L. & Elías-Arnanz, M. A new facet of vitamin B12: gene regulation by cobalamin-based photoreceptors. Annual review of biochemistry 86, 485–514 (2017).

40 Chatelle, C. et al. A Green-Light-Responsive System for the Control of Transgene Expression in Mammalian and Plant Cells. ACS Synth Biol 7, 1349–1358, doi:10.1021/acssynbio.7b00450 (2018).

41 Yeager, M. & Nicholson, B. J. Structure of gap junction intercellular channels. Current opinion in structural biology 6, 183–192 (1996).

42 Beyer, E. C., Paul, D. L. & Goodenough, D. A. Connexin43: a protein from rat heart homologous to a gap junction protein from liver. The Journal of cell biology 105, 2621–2629 (1987).

43 Jordan, K. et al. Trafficking, assembly, and function of a connexin43-green fluorescent protein chimera in live mammalian cells. Molecular biology of the cell 10, 2033–2050 (1999).

44 Campbell, R. E. et al. A monomeric red fluorescent protein. Proceedings of the National Academy of Sciences 99, 7877–7882 (2002).

45 Reed, K. et al. Molecular cloning and functional expression of human connexin37, an endothelial cell gap junction protein. The Journal of clinical investigation 91, 997–1004 (1993).

46 Chen, H., Li, Y. X., Wong, R. S., Esseltine, J. L. & Bai, D. Genetically engineered human embryonic kidney cells as a novel vehicle for dual patch clamp study of human gap junction channels. Biochemical Journal 481, 741–758 (2024).

47 Jassim, A., Aoyama, H., Willy, G. Y., Chen, H. & Bai, D. Engineered Cx40 variants increased docking and function of heterotypic Cx40/Cx43 gap junction channels. Journal of Molecular and Cellular Cardiology 90, 11–20 (2016).

48 Willy, G. Y. et al. Junctional delay, frequency, and direction-dependent uncoupling of human heterotypic Cx45/Cx43 gap junction channels. Journal of molecular and cellular cardiology 111, 17–26 (2017).

49 Czyż, J., Irmer, U., Schulz, G., Mindermann, A. & Hülser, D. F. Gap-junctional coupling measured by flow cytometry. Experimental Cell Research 255, 40–46 (2000).

50 Juul, M. H., Rivedal, E., Stokke, T. & Sanner, T. Quantitative determination of gap junction intercellular communication using flow cytometric measurement of fluorescent dye transfer. Cell adhesion and communication 7, 501–512 (2000).

51 Nielsen, M. J., Rasmussen, M. R., Andersen, C. B., Nexø, E. & Moestrup, S. K. Vitamin B12 transport from food to the body’s cells—a sophisticated, multistep pathway. Nature reviews Gastroenterology & hepatology 9, 345–354 (2012).

52 Gick, G. G. et al. Cellular uptake of vitamin B12: Role and fate of TCblR/CD320, the transcobalamin receptor. Experimental Cell Research 396, 112256 (2020).

53 Yang, Z. et al. B12-induced reassembly of split photoreceptor protein enables photoresponsive hydrogels with tunable mechanics. Science Advances 8, eabm5482 (2022).

54 Nutrients, S. o. U. R. L. o., Intakes, S. C. o. t. S. E. o. D. R., Folate, i. P. o., Vitamins, O. B. & Choline. Dietary reference intakes for thiamin, riboflavin, niacin, vitamin B6, folate, vitamin B12, pantothenic acid, biotin, and choline. (2000).

55 Wu, J. & Chen, Z. J. Innate immune sensing and signaling of cytosolic nucleic acids. Annual review of immunology 32, 461–488 (2014).

56 Ergun, S. L., Fernandez, D., Weiss, T. M. & Li, L. STING polymer structure reveals mechanisms for activation, hyperactivation, and inhibition. Cell 178, 290–301. e210 (2019).

57 Wei, X., et al. LL-37 transports immunoreactive cGAMP to activate STING signaling and enhance interferon-mediated host antiviral immunity. Cell Reports 39 (2022).

58 Xu, Y. et al. Hybrid indicators for fast and sensitive voltage imaging. Angewandte Chemie 130, 4013–4017 (2018).

59 Zou, P. et al. Bright and fast multicoloured voltage reporters via electrochromic FRET. Nature communications 5, 4625 (2014).

60 Choi, E. J., Palacios-Prado, N., Sáez, J. C. & Lee, J. Identification of Cx45 as a major component of GJs in HeLa cells. Biomolecules 10, 1–14 (2020).

61 Phelan, P. Innexins: members of an evolutionarily conserved family of gap-junction proteins. Biochimica et Biophysica Acta (BBA)-Biomembranes 1711, 225–245 (2005).

62 Kirilly, D., Wang, S. & Xie, T. Self-maintained escort cells form a germline stem cell differentiation niche. Development 138, 5087–5097 (2011).

63 Zhu, C.-H. & Xie, T. Clonal expansion of ovarian germline stem cells during niche formation in Drosophila. (2003).

64 Sahai-Hernandez, P. & Nystul, T. G. A dynamic population of stromal cells contributes to the follicle stem cell niche in the Drosophila ovary. Development 140, 4490–4498 (2013).

65 Van Doren, M., Williamson, A. L. & Lehmann, R. Regulation of zygotic gene expression in Drosophila primordial germ cells. Current biology 8, 243–246 (1998).

66 Osterwalder, T., Yoon, K. S., White, B. H. & Keshishian, H. A conditional tissue-specific transgene expression system using inducible GAL4. Proceedings of the National Academy of Sciences 98, 12596–12601 (2001).

67 Hendi, A. et al. Channel-independent function of UNC-9/Innexin in spatial arrangement of GABAergic synapses in C. elegans. Elife 11, e80555 (2022).

68 Das, M., Cheng, D., Matzat, T. & Auld, V. J. Innexin-mediated adhesion between glia is required for axon ensheathment in the peripheral nervous system. Journal of Neuroscience 43, 2260–2276 (2023).

69 Miao, G., Godt, D. & Montell, D. J. Integration of migratory cells into a new site in vivo requires channel-independent functions of innexins on microtubules. Developmental cell 54, 501–515. e509 (2020).

70 Goodenough, D. A. & Paul, D. L. Beyond the gap: functions of unpaired connexon channels. Nature reviews Molecular cell biology 4, 285–295 (2003).

71 Sjöstrand, F., Andersson-Cedergren, E. & Dewey, M. The ultrastructure of the intercalated discs of frog, mouse and guinea pig cardiac muscle. Journal of ultrastructure research 1, 271–287 (1958).

72 Kyung, T. et al. Optogenetic control of endogenous Ca2+ channels in vivo. Nature biotechnology 33, 1092–1096 (2015).

73 Banaszynski, L. A., Liu, C. W. & Wandless, T. J. Characterization of the FKBP⊙ Rapamycin⊙ FRB ternary complex. Journal of the American Chemical Society 127, 4715–4721 (2005).

74 Zhang, Q. et al. Visualizing dynamics of cell signaling in vivo with a phase separation-based kinase reporter. Molecular cell 69, 334–346. e334 (2018).

75 Li, J., Kim, S. G. & Blenis, J. Rapamycin: one drug, many effects. Cell metabolism 19, 373–379 (2014).

76 Laplante, M. & Sabatini, D. M. mTOR signaling in growth control and disease. cell 149, 274–293 (2012).

77 Smyth, J. W. & Shaw, R. M. Autoregulation of connexin43 gap junction formation by internally translated isoforms. Cell reports 5, 611–618 (2013).

78 Roth, W. & Mohamadzadeh, M. Vitamin B12 and gut-brain homeostasis in the pathophysiology of ischemic stroke. EBioMedicine 73 (2021).

79 Wei, C. J., Xu, X. & Lo, C. W. Connexins and cell signaling in development and disease. Annu Rev Cell Dev Biol 20, 811–838, doi:10.1146/annurev.cellbio.19.111301.144309 (2004).

80 Goodenough, D. A. & Paul, D. L. Gap junctions. Cold Spring Harb Perspect Biol 1, a002576, doi:10.1101/cshperspect.a002576 (2009).

