## Supplementary Material for "Controllable Gap Junctions by Vitamin B_12_ and Light"

**Affiliations:**

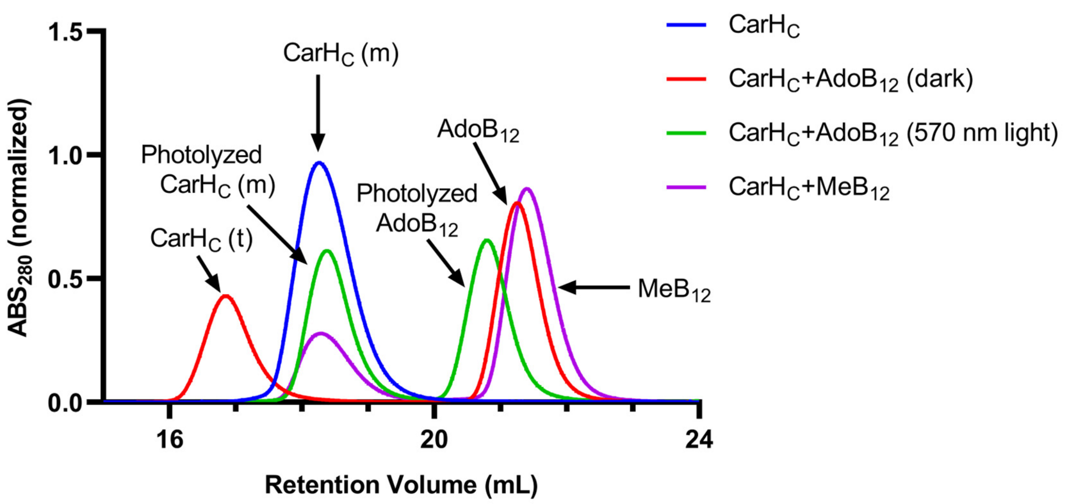


**Supplementary Figure 1 | SEC analyses of CarH$_C$ with addition of AdoB$_{12}$, MeB$_{12}$.** **"m" and "t" denote "monomer" and "tetramer", respectively.**

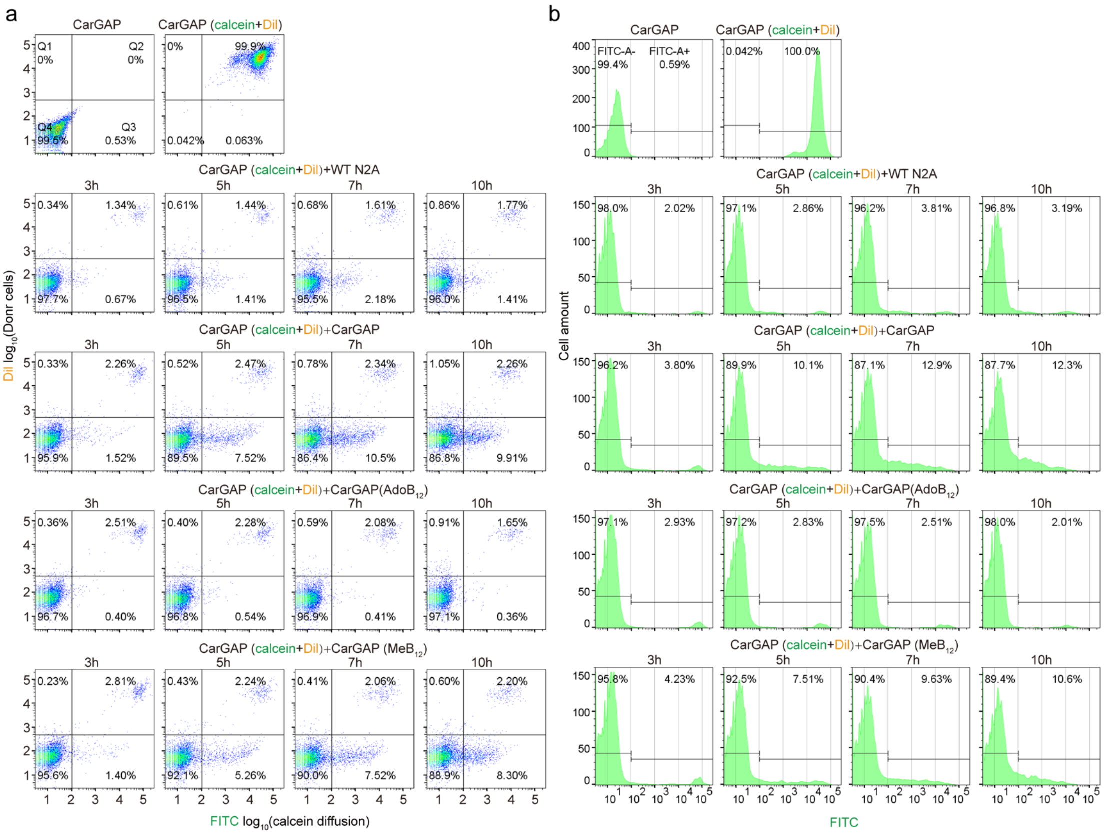


**Supplementary Figure 2 | Time-course analysis of gap junction reformation between co-cultured cells by flow cytometry. a**, Flow cytometry plots showing four quadrants (Q1-Q4). Donor cells (Dil+/calcein+, Q2) were co-cultured with recipient cells at a 1:40 ratio. Calcein transfer to recipient cells (Q3) was measured at 3, 5, 7, and 10 h post-co-culturing. At 3 h, minimal transfer was observed in: WT N2A (0.67%), CarGAP N2A (1.52%), AdoB12-treated CarGAP N2A (0.40%), and MeB12-treated CarGAP N2A (1.40%). By 5 h, significant differences emerged: WT N2A (1.41%), CarGAP N2A (7.52%), AdoB12-treated CarGAP N2A (0.54%), and MeB12-treated CarGAP N2A (5.26%), establishing 5 h as the optimal timepoint for functional assessment of GJCs. **b**, Histograms showing calcein distribution between donor and recipient cell populations at each timepoint.

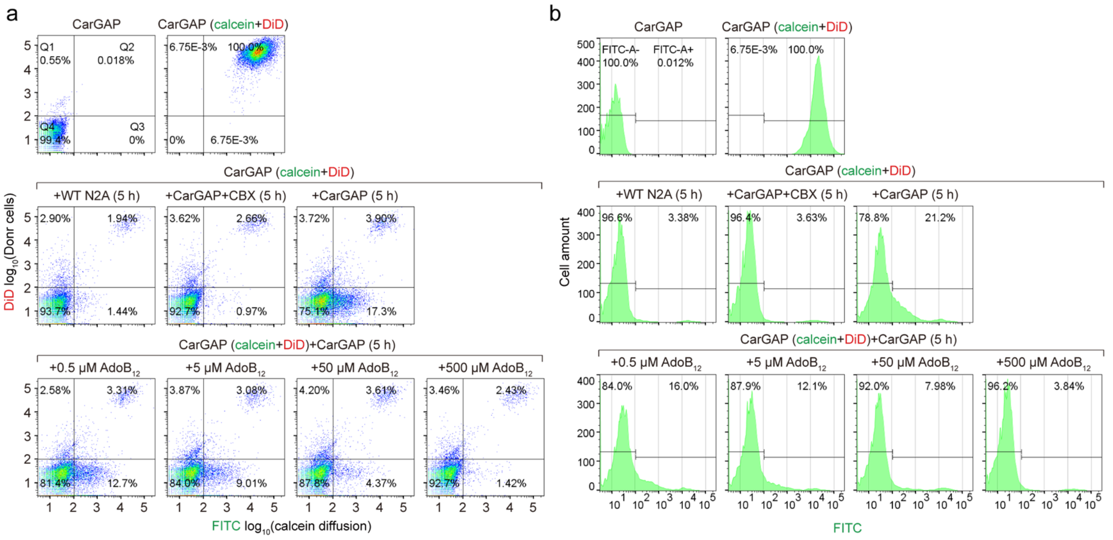


**Supplementary Figure 3 | Influence of varied amounts of $AdoB_{12}$ on CarGAP. a,** Flow cytometry analysis of calcein transfer from DiD+/calcein+ donor cells (Q2) to recipient cells (Q3) at varying $AdoB_{12}$ concentrations (0, 0.5, 5, 50, and 500 µM). The donor-to-recipient ratio was ~1:40. Untreated CarGAP N2A cells exhibited 17.3% calcein transfer, which decreased dose-dependently to 12.7% (0.5 µM $AdoB_{12}$), 9.01% (5 µM $AdoB_{12}$), 4.37% (50 µM $AdoB_{12}$), and 1.42% (500 µM $AdoB_{12}$) which is comparable to 0.97% (100 µM CBX), demonstrating complete inhibition at 500 µM $AdoB_{12}$. (b) **b**, Corresponding histogram profiles of calcein distribution across experimental conditions.

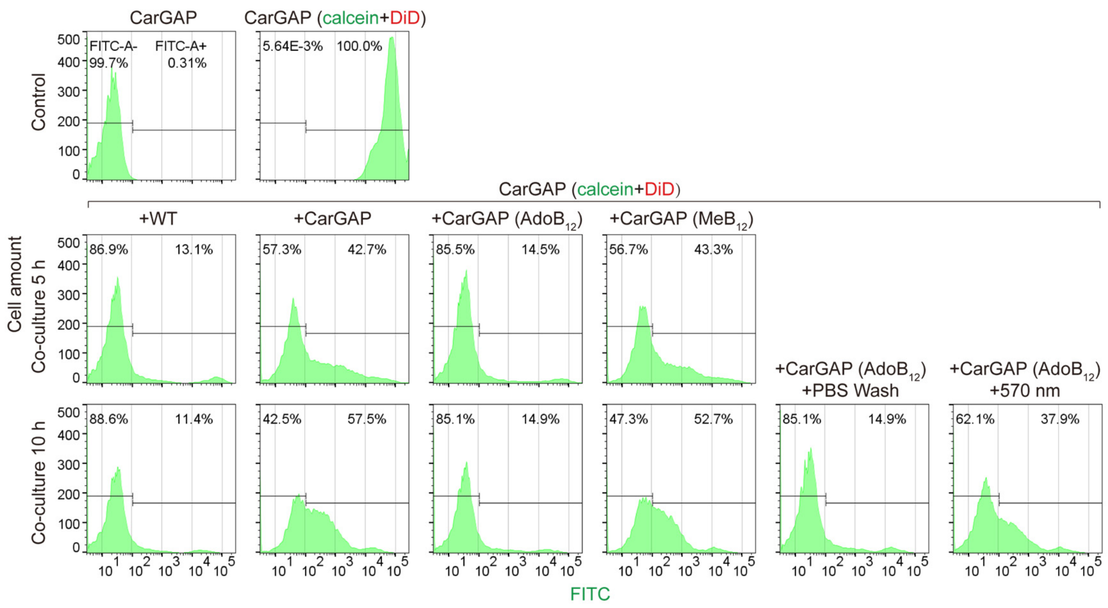


**Supplementary Figure 4 | Flow cytometric analysis of calcein transfer between N2A cells.** Histograms show calcein distribution profiles assessing CarGAP functionality. Using a FITC intensity threshold of $10^2$, we distinguished between autofluorescent (FITC-A-) and calcein-positive (FITC-A+) cell populations. The integrated percentages of FITC-A- and FITC-A+ quantify baseline autofluorescence and calcein transfer efficiency, respectively.

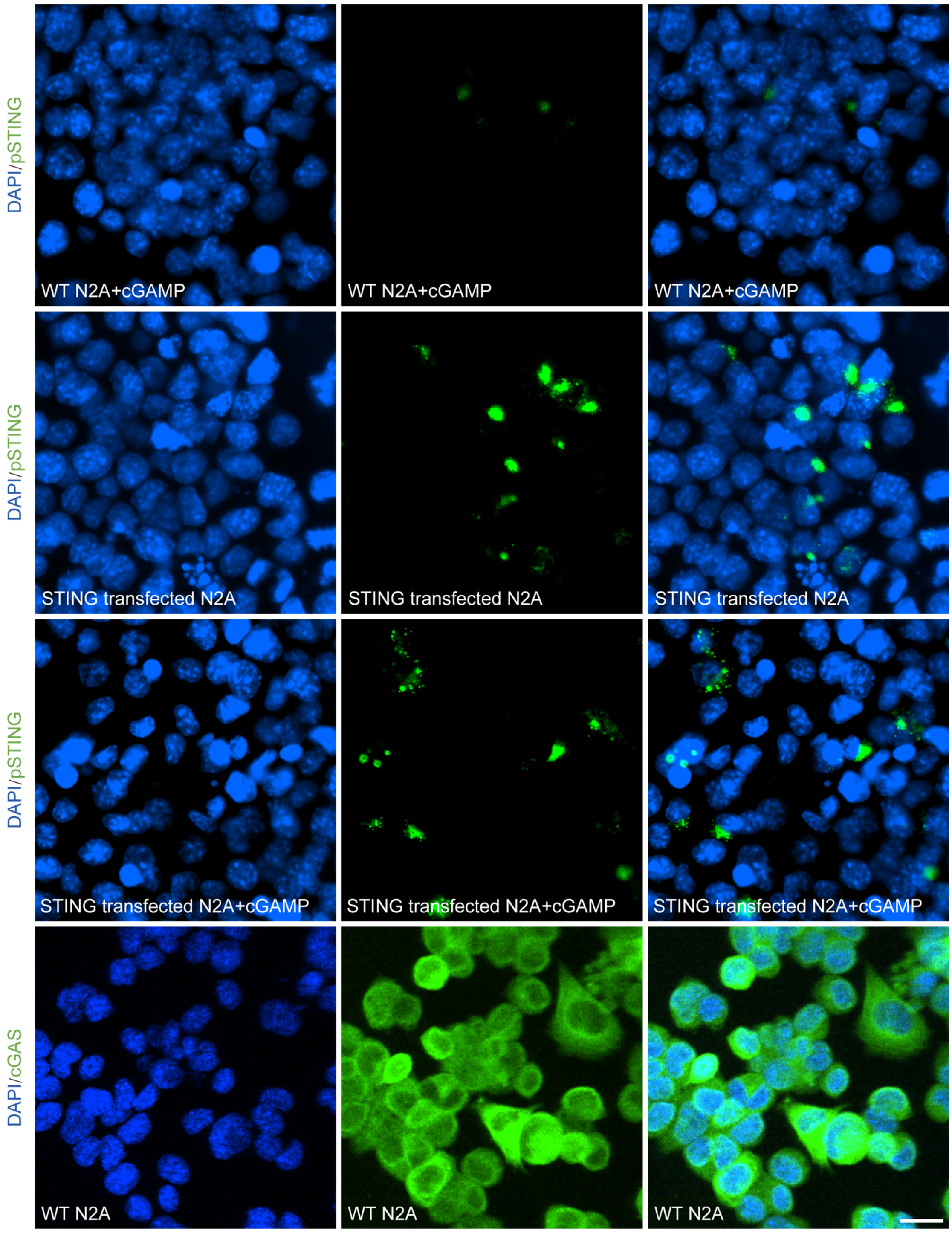


**Supplementary Figure 5 | Immunofluorescence images showing few endogenous STING and abundant endogenous cGAS in wild-type (WT) N2A cells.** The phosphorylated STING proteins were immunostained using a rabbit anti-pSTING antibody (Cell Signaling, #62912). The cGAS proteins were immunostained using a rabbit anti-cGAS antibody (Cell Signaling, #79978). Scale bar: 20 µm.

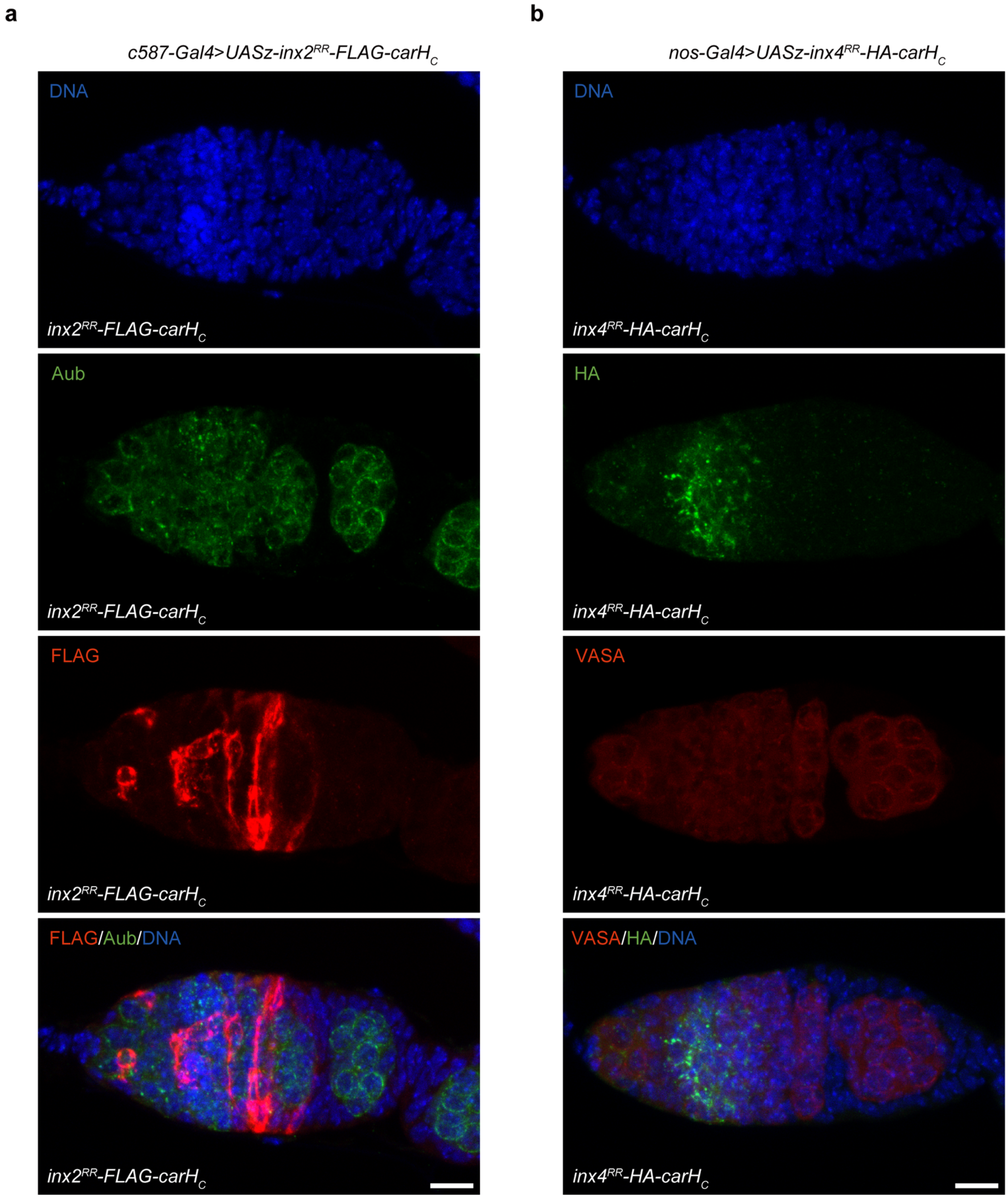


**Supplementary Figure 6 | Expression and localization of Inx2-FLAG-CarH$_C$ in somatic cells and Zpg-HA-CarH$_C$ in germ cells within *Drosophila* germaria. a**, Expression of Inx2-FLAG-CarH$_C$ in escort cells was driven by c587-Gal4, with the protein localizing specifically to the membrane edge of these cells. DNA is stained by DAPI (blue channel). Germline is stained by anti-Aub antibody (green channel). Inx2-CarH$_C$ is stained by anti-FLAG antibody (red channel). Scale bar: 10 µm. **b**, Expression of Zpg-HA-CarH$_C$ in germ cells was driven by nos-Gal4, with the protein localizing specifically to the membrane edge of these cells. DNA is stained by DAPI (blue channel). Zpg-CarH$_C$ is stained by anti-HA antibody (green channel). Germline is stained by anti-Vasa antibody (red channel). Scale bar: 10 µm.

a

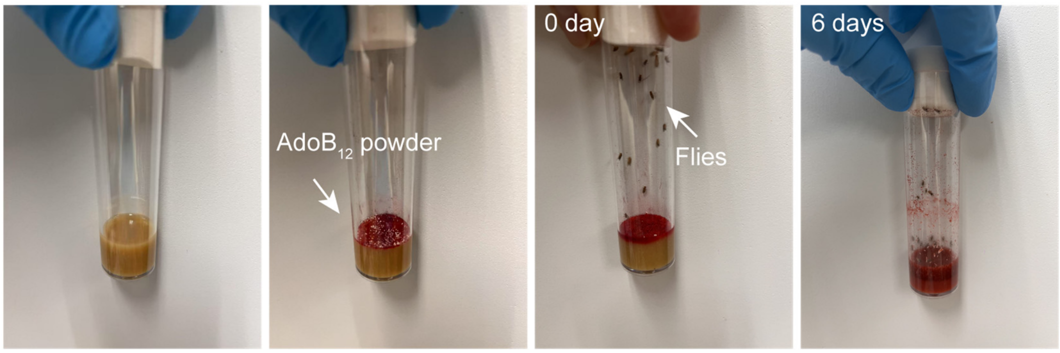


b

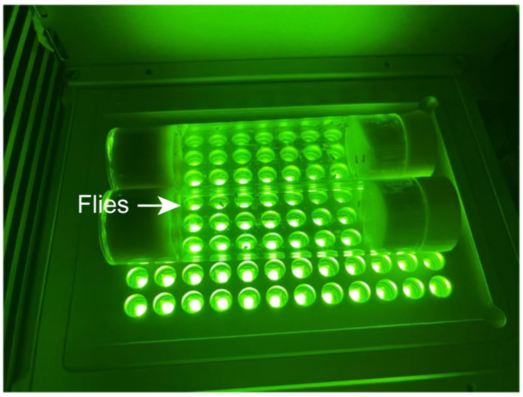


**Supplementary Figure 7 | AdoB$_{12}$ administration and green light illumination setup for *Drosophila*.** **a**, Flies were cultured in a tube containing *Drosophila* media supplemented with yeast extract and AdoB$_{12}$ (0.01 g per tube), evenly distributed on the food surface. **b**, Green light illumination apparatus showing cold LED placement below the culture tubes ($\lambda$ = 570 nm, 80 mW/cm$^2$). Light intensity was calibrated by voltage adjustment and verified using an optical power meter according to manufacturer specifications.

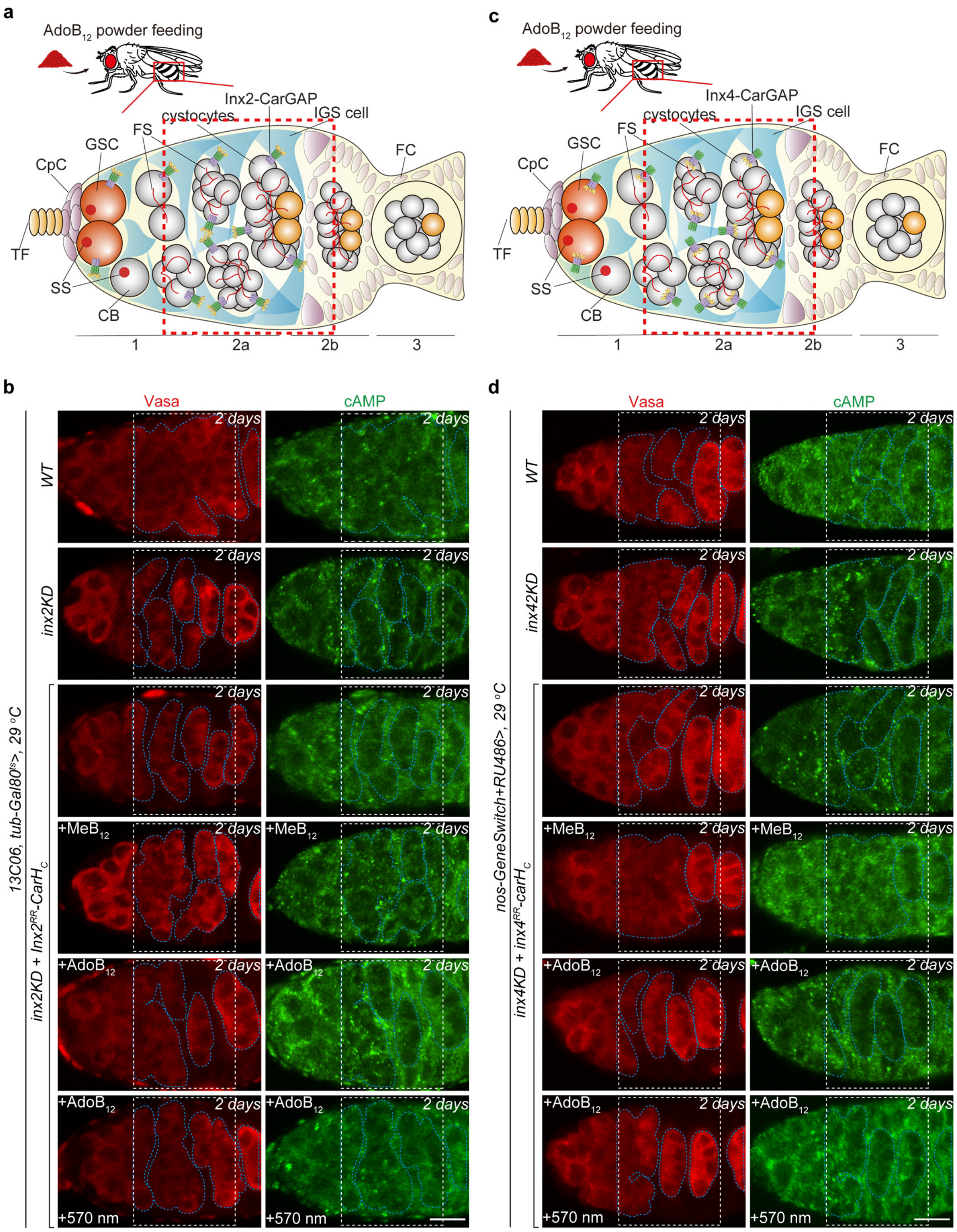


**Supplementary Figure 8 | Detailed methods for quantifying cAMP fluorescence intensity in IGS/germline cells.** Representative images showing the quantification regions (white dotted rectangle, S1) spanning stage 2a and proximal stage 2b of germarium development in Inx2-CarGAP flies (**a**) and Inx4-CarGAP flies (**b**). Germline boundary demarcation (blue dotted lines, S2) was determined based on Vasa staining. cAMP fluorescence intensity ratios were calculated as:

$$\frac{\mathrm{IGS}}{\mathrm{Germline}} = \frac{\mathrm{S1} - \mathrm{S2}}{\mathrm{S2}}$$

**Supplementary Table 1: DNA sequence information**

| *Cx43*-*Linker*-*FLAG*-*carHc* |
| --- |
| GCCACCATGGGTGACTGGAGCGCCTTAGGCAAACTCCTTGACAAGGTTCA<br>AGCCTACTCAACTGCTGGAGGGAAGGTGTGGCTGTCAGTACTTTTCATTTT<br>CCGAATCCTGCTGCTGGGGACAGCGGTTGAGTCAGCCTGGGGAGATGAGC<br>AGTCTGCCTTTCGTTGTAACACTCAGCAACCTGGTTGTGAAAATGTCTGCT<br>ATGACAAGTCTTTCCCAATCTCTCATGTGCGCTTCTGGGTCCTGCAGATCA<br>TATTTGTGTCTGTACCCACACTCTTGTACCTGGCTCATGTGTTCTATGTGAT<br>GCGAAAGGAAGAGAAACTGAACAAGAAAGAGGAAGAACTCAAGGTTGCC<br>CAAACTGATGGTGTCAATGTGGACATGCACTTGAAGCAGATTGAGATAAA<br>GAAGTTCAAGTACGGTATTGAAGAGCATGGTAAGGTGAAAATGCGAGGG<br>GGGTTGCTGCGAACCTACATCATCAGTATCCTCTTCAAGTCTATCTTTGAG<br>GTGGCCTTCTTGCTGATCCAGTGGTACATCTATGGATTCAGCTTGAGTGCT<br>GTTTACACTTGCAAAAGAGATCCCTGCCCACATCAGGTGGACTGTTTCCTC<br>TCTCGCCCCACGGAGAAAACCATCTTCATCATCTTCATGCTGGTGGTGTCC<br>TTGGTGTCCCTGGCCTTGAATATCATTGAACTCTTCTATGTTTTCTTCAAGG<br>GCGTTAAGGATCGGGTTAAGGGAAAGAGCGACCCTTACCATGCGACCAGT<br>GGTGCGCTGAGCCCTGCCAAAGACTGTGGGTCTCAAAAATATGCTTATTT<br>CAATGGCTGCTCCTCACCAACCGCTCCCCTCTCGCCTATGTCTCCTCCTGG<br>GTACAAGCTGGTTACTGGCGACAGAAACAATTCTTCTTGCCGCAATTACA<br>ACAAGCAAGCAAGTGAGCAAAACTGGGCTAATTACAGTGCAGAACAAAA<br>TCGAATGGGGCAGGCGGGAAGCACCATCTCTAACTCCCATGCACAGCCTT<br>TTGATTTCCCCGATGATAACCAGAATTCTAAAAAACTAGCTGCTGGACAT<br>GAATTACAGCCACTAGCCATTGTGGACCAGCGACCTTCAAGCAGAGCCAG<br>CAGTCGTGCCAGCAGCAGACCTCGGCCTGATGACCTGGAGATCGGCGGCA<br>GTGGAGGAAGCGACTATAAAGATGATGACGACAAGCCAGAAGATCTTGGC<br>ACTGGCCTTCTTGAAGCACTGCTCCGGGGAGATCTTGCGGGCGCTGAGGC<br>GTTGTTCAGACGAGGGCTTAGATTTTGGGGCCCAGAAGGCGTACTTGAAC<br>ACCTGCTCTTGCCTGTCCTTCGGGAAGTGGGAGAAGCATGGCACCGAGGG<br>GAGATCGGAGTAGCTGAGGAGCATCTGGCCTCTACATTTCTCCGCGCAAGA<br>TTGCAAGAACTCTTGGACTTGGCAGGCTTCCCTCCTGGCCCACCGGTACTG<br>GTAACGACACCCCCGGGCGAGCGCCACGAGATTGGCGCCATGTTGGCGGC<br>ATACCACCTGAGACGAAAAGGTGTCCCTGCGCTTTACCTTGGACCAGATAC<br>ACCACTTCCTGATTTGCGAGCATTGGCTAGAAGACTCGGTGCAGGGGCTGT<br>TGTGTTGTCAGCCGTACTGTCTGAACCACTCAGAGCTCTCCCGGATGGGGC<br>GCTGAAAGATTTGGCGCCTCGGGTCTTTTTGGGGGGTCAGGGCGCTGGCC<br>CTGAGGAGGCGAGGCGCCTTGGGGCCGAATATATGGAGGACCTGAAGGGT<br>CTTGCTGAAGCACTGTGGCTGCCTCGAGGACCTGAAAAAGAGGCCATA |
| *cGAS* |
| GCCGCCACCATGCAACCGTGGCATGGGAAAGCAATGCAGCGCGCTTCCGA<br>GGCTGGGGCTACTGCTCCTAAAGCGAGCGCCCGCAACGCGAGAGGTGCGC<br>CAATGGATCCTACGGAAAGTCCAGCCGCTCCCGAAGCAGCGCTGCCAAAA<br>GCGGGGAAGTTTGGGCCTGCCCGCAAGAGTGGTTCACGGCAAAAAAAAA |

| |
|---|
| GCGCACCAGATACGCAGGAACGCCCACCAGTAAGGGCTACCGGCGCGCGA<br>GCTAAGAAAGCACCCCAGCGCGCTCAAGACACTCAGCCCTCTGATGCAAC<br>GTCAGCTCCTGGTGCGGAGGGTTTGGAGCCGCCTGCGGCTAGGGAACCGG<br>CACTCTCACGAGCCGGCTCTTGTCGGCAGAGAGGCGCACGGTGTTCCACG<br>AAACCACGACCTCCCCCTGGGCCATGGGATGTGCCTTCACCTGGCCTTCCC<br>GTCTCCGCGCCAATTTTGGTTCGGAGGGATGCTGCGCCGGGGGCGTCCAA<br>ATTGCGCGCGGTCTTGGAAAAGCTCAAGCTGTCTCGCGATGACATCAGCAC<br>AGCCGCAGGAATGGTGAAGGGCGTAGTAGATCACCTTCTGCTCAGACTGA<br>AATGTGACAGCGCATTTAGGGGGGTCGGGCTCCTCAATACAGGGTCCTATT<br>ATGAGCACGTAAAAATATCAGCACCAAATGAGTTTGACGTAATGTTCAAAC<br>TGGAAGTTCCGCGCATCCAGTTGGAGGAGTATAGTAATACACGAGCCTACT<br>ATTTTGTAAAGTTCAAACGCAACCCCAAGGAGAACCCGTTGTCACAATTCC<br>TCGAGGGGGAAATACTCAGTGCCTCTAAGATGTTGTCAAAGTTCCGGAAG<br>ATCATAAAAGAGGAGATAAACGATATTAAAGACACTGACGTAATAATGAAA<br>AGAAAACGGGGAGGTTCACCGGCCGTTACTCTCTTGATTAGTGAAAAAATT<br>TCTGTGGACATCACCCTCGCACTGGAGTCCAAGTCTTCCTGGCCTGCAAGC<br>ACCCAGGAAGGTCTCCGAATTCAGAATTGGTTGAGTGCCAAAGTTAGAAA<br>ACAGTTGCGACTCAAACCTTTCTACTTGGTCCCAAAACACGCCAAGGAGG<br>GAAATGGCTTTCAAGAGGAGACTTGGCGCCTCAGTTTCAGTCACATAGAG<br>AAAGAGATTCTTAATAACCACGGTAAAAGTAAAACCTGTTGCGAAAATAAA<br>GAAGAGAAATGTTGCAGGAAAGACTGTCTCAAGCTGATGAAATACCTTCT<br>GGAGCAGCTCAAGGAACGATTCAAAGATAAGAAACACCTGGACAAGTTCT<br>CTAGCTACCACGTAAAAACTGCCTTCTTTCATGTATGTACGCAGAACCCCCA<br>GGATAGTCAGTGGGACAGGAAGGATCTCGGCCTGTGTTTCGATAATTGTGT<br>CACTTATTTTCTCCAATGCCTCCGAACTGAGAAGCTGGAAAACTACTTCATA<br>CCTGAGTTCAATCTTTTCTCAAGCAACTTGATAGATAAACGAAGCAAAGAG<br>TTTTTGACAAAGCAAATTGAATATGAAAGGAACAATGAATTCCCGGTTTTT<br>GATGAGTTC |
| *STING* |
| GCCGCCACCATGCCACACTCTTCCTTGCATCCATCCATACCTTGCCCTAGAG<br>GACATGGAGCTCAGAAAGCAGCCTTGGTACTTCTCTCAGCTTGTCTGGTGA<br>CACTCTGGGGCCTCGGCGAGCCCCCGAGCACACTCTCCGATATCTTGTTT<br>TGCATTTGGCTAGCTTGCAACTCGGTTTGCTTCTTAATGGGGTGTGCTCACT<br>GGCGGAGGAGCTGCGCCACATCCACTCACGGTATAGAGGTTCCTATTGGCG<br>GACTGTTCGCGCTTGTCTGGGGTGCCCATTGAGGAGAGGAGCCCTCCTTCT<br>TTTGTCCATTTATTTCTATTACAGTCTTCCAAATGCGGTTGGACCTCCATTCA<br>CTTGGATGTTGGCCCTCCTCGGTCTTTCTCAAGCCCTGAACATCCTCCTGGG<br>GCTCAAAGGCCTCGCGCCTGCTGAAATTAGTGCAGTGTGCGAAAAGGGAA<br>ATTTTAATGTGGCACACGGCTTGGCTTGGTCTTATTATATCGGGTACCTCCGA<br>CTGATACTGCCCGAATTGCAGGCCCGCATTAGGACGTACAATCAACATTATA<br>ACAATCTGCTGAGGGGTGCTGTAAGTCAACGACTCTACATCCTGTTGCCCT<br>TGGACTGCGGAGTCCCCGATAATCTCTCTATGGCAGATCCAAATATAAGGTT<br>CCTTGACAAACTTCCCCAACAGACTGGTGACCGCGCCGGAATCAAGGACA<br>GGGTTTATTCTAACTCCATCTATGAACTCCTTGAGAACGGACAGCGGGCCG |

| |
|---|
| GTACCTGCGTCCTTGAGTACGCCACTCCTCTCCAAACCCTGTTCGCTATGTC<br>CCAGTACTCACAGGCAGGATTCAGTCGAGAAGATCGGTTGGAGCAGGCCA<br>AGCTTTTCTGTCGGACTCTGGAAGACATACTTGCAGATGCCCCAGAGTCAC<br>AAAACAATTGCCGGTTGATAGCCTATCAGGAACCCGCCGATGACTCTAGTT<br>TTAGCCTGTCACAAGAGGTACTGCGACATCTCCGGCAGGAAGAGAAAGAA<br>GAAGTGACTGTTGGTTCCCTGAAAACTTCCGCAGTCCCTAGCACTAGCACG<br>ATGTCTCAAGAGCCCGAATTGCTGATAAGCGGAATGGAAAAGCCACTCCCT<br>CTCCGAACGGACTTTAGT |
| *STING-egfp* |
| GCCGCCACCATGCCACACTCTTCCTTGCATCCATCCATACCTTGCCCTAGAG<br>GACATGGAGCTCAGAAAGCAGCCTTGGTACTTCTCTCAGCTTGTCTGGTGA<br>CACTCTGGGGCCTCGGCGAGCCCCCGAGCACACTCTCCGATATCTTGTTT<br>TGCATTTGGCTAGCTTGCAACTCGGTTTGCTTCTTAATGGGGTGTGCTCACT<br>GGCGGAGGAGCTGCGCCACATCCACTCACGGTATAGAGGTTCCTATTGGCG<br>GACTGTTCGCGCTTGTCTGGGGTGCCCATTGAGGAGAGGAGCCCTCCTTCT<br>TTTGTCCATTTATTTCTATTACAGTCTTCCAAATGCGGTTGGACCTCCATTCA<br>CTTGGATGTTGGCCCTCCTCGGTCTTTCTCAAGCCCTGAACATCCTCCTGGG<br>GCTCAAAGGCCTCGCGCCTGCTGAAATTAGTGCAGTGTGCGAAAAGGGAA<br>ATTTTAATGTGGCACACGGCTTGGCTTGGTCTTATTATATCGGGTACCTCCGA<br>CTGATACTGCCCGAATTGCAGGCCCGCATTAGGACGTACAATCAACATTATA<br>ACAATCTGCTGAGGGGTGCTGTAAGTCAACGACTCTACATCCTGTTGCCCT<br>TGGACTGCGGAGTCCCCGATAATCTCTCTATGGCAGATCCAAATATAAGGTT<br>CCTTGACAAACTTCCCCAACAGACTGGTGACCGCGCCGGAATCAAGGACA<br>GGGTTTATTCTAACTCCATCTATGAACTCCTTGAGAACGGACAGCGGGCCG<br>GTACCTGCGTCCTTGAGTACGCCACTCCTCTCCAAACCCTGTTCGCTATGTC<br>CCAGTACTCACAGGCAGGATTCAGTCGAGAAGATCGGTTGGAGCAGGCCA<br>AGCTTTTCTGTCGGACTCTGGAAGACATACTTGCAGATGCCCCAGAGTCAC<br>AAAACAATTGCCGGTTGATAGCCTATCAGGAACCCGCCGATGACTCTAGTT<br>TTAGCCTGTCACAAGAGGTACTGCGACATCTCCGGCAGGAAGAGAAAGAA<br>GAAGTGACTGTTGGTTCCCTGAAAACTTCCGCAGTCCCTAGCACTAGCACG<br>ATGTCTCAAGAGCCCGAATTGCTGATAAGCGGAATGGAAAAGCCACTCCCT<br>CTCCGAACGGACTTTAGTGGTGGGGGCGGTAGCGTCAGTAAAGGAGAGGA<br>ACTGTTCACTGGTGTAGTACCAATTTTGGTTGAGCTTGATGGTGACGTTAAT<br>GGACACAAATTTTCTGTCTCAGGTGAGGGCGAGGGAGATGCTACCTACGGT<br>AAATTGACTTTGAAATTCATTTGTACGACAGGAAAGTTGCCCGTCCCATGG<br>CCAACACTGGTAACTACGCTGACCTATGGTGTCCAATGTTTCTCCCGGTACC<br>CCGACCACATGAAACAACACGATTTTTTCAAGTCTGCAATGCCAGAAGGGT<br>ATGTTCAGGAAAGAACTATTTTTTTCAAAGACGATGGTAACTACAAAACAA<br>GAGCGGAGGTAAAGTTTGAAGGAGACACGCTTGTTAATCGCATAGAGCTG<br>AAGGGGATCGACTTCAAAGAAGATGGAAACATTCTTGGGCATAAGTTGGA<br>GTATAATTACAATAGTCACAATGTCTATATAATGGCGGACAAACAGAAAAAT<br>GGAATTAAAGTAAACTTTAAAATCCGCCATAACATTGAAGACGGTAGCGTT<br>CAACTTGCGGACCACTATCAGCAAAACACCCCTATAGGTGACGGCCCCGTA<br>TTGTTGCCGGACAATCATTATCTTTCCACACAAAGCAAGTTGTCTAAAGATC |

| |
|---|
| CCAACGAAAAGAGAGATCACATGGTACTTCTCGAGTTTGTCACGGCTGCG<br>GGGATTACCCTGGGTATGGACGAGCTTTACAAA |
| *inx2*$^{RR}$-*FLAG*-*carHc* |
| CACCATGTTTGATGTCTTTGGGTCCGTCAAGGGCCTGCTGAAGATCGACCA<br>GGTGTGCATCGACAACAATGTCTTTCGCATGCACTACAAGGCCACGGTGAT<br>CATACTGATCGCCTTCTCCCTCCTGGTGACCTCGCGCCAATACATCGGTGAC<br>CCCATCGATTGTATTGTGGACGAGATCCCACTGGGCGTGATGGACACCTAC<br>TGCTGGATCTACTCTACGTTTACCGTGCCAGAGAGGCTAACGGGCATCACC<br>GGACGCGATGTGGTGCAGCCCGGCGTGGGCTCCCATGTGGAGGGCGAGGA<br>CGAGGTGAAGTACCACAAGTACTACCAGTGGGTGTGCTTCGTCCTCTTCTT<br>CCAGGCCATCCTGTTCTACGTACCGCGCTATCTGTGGAAGTCCTGGGAAGG<br>CGGACGCCTCAAGATGCTGGTCATGGATCTCAAAAGTCCCATCGTTAATGA<br>CGAGTGCAAGAACGATCGCAAAAAGATCCTGGTCGACTACTTCATTGGCA<br>ACCTGAACCGCCACAATTTCTACGCCTTCCGATTCTTCGTGTGCGAAGCCC<br>TGAACTTTGTGAATGTGATTGGACAGATCTACTTTGTGGACTTCTTCCTCGA<br>CGGCGAGTTCAGCACCTACGGCAGTGATGTCTTGAAGTTCACTGAGCTGG<br>AGCCGGATGAGCGCATTGATCCCATGGCGCGGGTCTTTCCGAAGGTCACCA<br>AATGCACGTTCCACAAATACGGTCCATCCGGCAGTGTGCAGACCCACGACG<br>GTCTGTGCGTGCTGCCCCTGAACATTGTCAACGAAAAGATCTACGTGTTCC<br>TGTGGTTCTGGTTCATCATCCTGAGCATCATGTCGGGAATATCGCTTATCTAC<br>AGAATCGCTGTTGTGGCGGGTCCCAAGCTCCGCCATCTCCTACTCCGAGCC<br>CGTTCCCGTTTGGCTGAAAGCGAGGAGGTCGAACTGGTGGCCAACAAGTG<br>CAACATCGGCGATTGGTTCCTGCTCTATCAGCTGGGCAAGAACATCGATCC<br>GCTCATCTACAAGGAGGTGATCTCGGACTTGTCCCGCGAAATGAGCGGCGA<br>TGAGCATAGCGCCCACAAGCGGCCCTTCGACGCCGATTACAAGGATGACG<br>ACGATAAGCCAGAAGATCTGGGCACCGGCCTGCTGGAAGCACTGCTGCGC<br>GGTGATCTGGCGGGCGCCGAAGCTCTGTTTCGTCGTGGCCTGCGTTTCTGG<br>GGCCCGGAAGGCGTTCTGGAGCACCTGCTGCTGCCGGTGCTGCGTGAAGT<br>GGGCGAAGCTTGGCACCGTGGTGAAATCGGCGTTGCAGAAGAACACCTGG<br>CGAGCACCTTCCTGCGCGCGCGTCTGCAGGAGCTGCTGGACCTGGCAGGT<br>TTCCCGCCGGGTCCGCCGGTCCTGGTGACTACGCCGCCGGGCGAACGCCA<br>CGAAATCGGTGCGATGCTGGCGGCGTACCATCTGCGTCGTAAGGGCGTCCC<br>GGCGCTGTATCTGGGCCCGGATACTCCGCTGCCGGACCTGCGTGCACTGGC<br>GCGCCGCCTGGGTGCAGGCGCGGTCGTGCTGTCTGCTGTTCTGAGCGAAC<br>CGCTGCGTGCTCTGCCTGACGGTGCCCTGAAAGATCTGGCACCGCGTGTTT<br>TCCTGGGCGGCCAGGGCGCAGGCCCGGAAGAGGCACGCCGTCTGGGTGC<br>CGAATACATGGAAGACCTGAAAGGCCTGGCTGAAGCGCTGTGGCTGCCGC<br>GCGGTCCGGAAAAAGAAGCAATC |
| *Zpg*$^{RR}$-*HA*-*carHc* |
| CACCATGTACGCAGCTGTTAAACCGCTCTCCAAGTATCTGCAGTTCAAGTC<br>GGTGCACATCTACGATGCTATTTTCACGCTGCACTCCAAAGTCACAGTGGC<br>CCTGCTTTTGGCCTGCACATTTTTGCTTTCCTCGAAACAATATTTCGGCGAT<br>CCCATCCAGTGTTTTGGGGACAAGGATATGGACTACGTGCACGCCTTTTGT<br>TGGATCTACGGGGCCTATGTGAGCGACAATGTGACTGTGACGCCCCTAAGG |

AATGGAGCTGCACAGTGCCGACCAGATGCGGTTAGTAAGGTAGTGCCCCC
GGAAAATCGTAACTACATAACCTACTATCAGTGGGTCGTGCTGGTATTGCTG
CTCGAGTCGTTTGTGTTTTATATGCCAGCATTTCTCTGGAAGATTTGGGAGG
GCGGTCGGCTGAAGCATTTGTGCGATGATTTCCACAAGATGGCCGTTTGCA
AGGACAAGAGCAGGACCCATTTGCGCGTACTAGTCAACTACTTCTCCAGCG
ATTACAAAGAGACCCACTTCCGCTACTTCGTTAGCTACGTCTTCTGCGAAAT
ACTCAATTTGAGCATCAGTATTCTAAACTTCCTTCTGTTGGACGTATTTTTCG
GAGGTTTTTGGGGCCGCTACCGTAATGCCTTGCTGTCCCTTTACAATGGCGA
CTATAACCAGTGGAACATTATAACCATGGCAGTCTTCCCCAAGTGTGCCAA
GTGCGAGATGTACAAGGGCGGCCCCAGCGGTTCTTCCAACATATACGACTA
CCTTTGCCTGCTGCCCCTGAATATACTGAACGAGAAAATCTTCGCCTTCCTG
TGGATCTGGTTCATCCTTGTAGCAATGCTCATTTCCCTCAAGTTTCTGTACC
GCCTGGCCACCGTTTTGTATCCCGGAATGCGTCTTCAGTTGCTTCGTGCCAG
GGCACGTTTCATGCCCAAGAAGCACTTGCAGGTGGCATTGCGCAATTGCAG
TTTTGGCGACTGGTTCGTGCTGATGCGCGTGGGCAACAACATCAGCCCCGA
GCTATTCCGTAAGTTGCTGGAGGAGCTGTACGAAGCTCAGTCCTTGATCAA
AATACCGCCAGGAGCGGACAAGATCTACCCATACGATGTTCCAGATTACGC
TCCAGAAGATCTGGGCACCGGCCTGCTGGAAGCACTGCTGCGCGGTGATC
TGGCGGGCGCCGAAGCTCTGTTTCGTCGTGGCCTGCGTTTCTGGGGCCCGG
AAGGCGTTCTGGAGCACCTGCTGCTGCCGGTGCTGCGTGAAGTGGGCGAA
GCTTGGCACCGTGGTGAAATCGGCGTTGCAGAAGAACACCTGGCGAGCAC
CTTCCTGCGCGCGCGTCTGCAGGAGCTGCTGGACCTGGCAGGTTTCCCGCC
GGGTCCGCCGGTCCTGGTGACTACGCCGCCGGGCGAACGCCACGAAATCG
GTGCGATGCTGGCGGCGTACCATCTGCGTCGTAAGGGCGTCCCGGCGCTGT
ATCTGGGCCCGGATACTCCGCTGCCGGACCTGCGTGCACTGGCGCGCCGCC
TGGGTGCAGGCGCGGTCGTGCTGTCTGCTGTTCTGAGCGAACCGCTGCGT
GCTCTGCCTGACGGTGCCCTGAAAGATCTGGCACCGCGTGTTTTCCTGGGC
GGCCAGGGCGCAGGCCCGGAAGAGGCACGCCGTCTGGGTGCCGAATACAT
GGAAGACCTGAAAGGCCTGGCTGAAGCGCTGTGGCTGCCGCGCGGTCCGG
AAAAAGAAGCAATC
